# Organ-specific filtering by abiotic and biotic environmental factors shapes distinct yet overlapping microbial communities across *Lotus corniculatus* roots, shoots, flowers, and seeds

**DOI:** 10.1101/2025.04.25.650623

**Authors:** Katrina Lutap, Oliver Bossdorf, Eric Kemen

## Abstract

Plant microbiome assembly is modulated via filtering by the host plant and local environment as well as stochastic processes like microbial dispersal. *Lotus corniculatus* in natural populations that are continuously exposed to natural perturbations and microbial sources is an ideal plant model system to study the ecological processes that structure the distinct yet overlapping microbial communities in plant organs. We observed spatial and temporal variation in microbiomes associated with *L. corniculatus* roots, shoots, flowers, and seeds across seven grassland sites for four years. In this study we examined how abiotic and biotic factors in the local environment throughout multiple years contribute to the structure of microbiomes associated with *L. corniculatus* populations. We show that plant microbiomes are shaped by a set of environmental factors that are distinct to each plant compartment. These environmental factors either directly influence the plant microbiomes or by indirectly affecting them via other biotic factors. The environmental factors soil temperature seasonality, soil microbiome composition, air temperature seasonality, plant community richness, and grazing are found to influence the structure and microbial interactions in the plant organs, and are different in relationships with microbiomes with each compartment, possibly influencing dispersal decisions of microorganisms and consequently contribute in shaping distinct yet overlapping microbiomes across plant organs. *Burkholderia* and *Sulfuritalea*, plant-associated microbes that are highly correlated with environmental variables across all plant organs, respond to environmental variables differently depending on their organ microhabitat. This organ-dependent environmental perception is also observed in biomarker microbes in roots, shoots, and flowers, such as the rhizobial symbiont *Mesorhizobium*, leaf pathogen *Setosphaeria*, and necrotroph *Botryotinia*, respectively. Our knowledge about the organ-specific response of plant microbiomes to abiotic and biotic perturbations will equip us with a framework to understand and engineer plant microbiomes in the context of global climate change. The observed patterns on dispersal decisions or habitat choice based on organ-dependent environmental cues and microbial interactions in plant microbiomes also advance our insights on how beneficial microbes or pathogens survive and persist on specific plant microhabitat and environmental conditions.

## INTRODUCTION

Plants are host to diverse and dynamic microbial communities and oftentimes these associations provide reciprocal benefits. Plants equip microorganisms with habitat and resources for survival and growth, while the associated microorganisms are crucial in plant growth and nutrition, plant health and defense, or stress tolerance (1). Beyond these relationships, plant microbiomes have the potential to be used as tools for manipulation and management of plants with improved health and productivity. To leverage such potential, an in depth understanding of the assembly processes that shape plant community structure and interactions is important. Current research has established a spatially and temporally dynamic microbial communities associated with plants. The observed spatial and temporal variations in these plant microbial communities can be attributed to factors such as geographic location, soil properties, plant genotype, plant compartment, seasonality, and plant age or developmental stages (2–7). Given their significant roles, understanding the mechanisms that drive community assembly and variation in plant microbiomes is essential.

One of the main deterministic factors that influence microbial community assembly is filtering by the host plant and local environment. Plant host filtering drives variation in diversity and composition of associated microbial communities when genetic and phenotypic variations impact survival and persistence of microorganisms in the host plant. Plant genomic loci associated with plant traits such as cell morphology, defense and immune responses, signaling pathways, and secondary metabolism affect variation of the associated microbial communities (8, 9). Morphological and chemical differences between plant organs also result in distinct microbial communities due to selective pressure by the unique microhabitat conditions (10–13). In addition to plant organ and genotype, filtering by abiotic and biotic factors in the local environment impacts the structure of plant microbial communities. Factors such as climate (*i.e.* temperature, precipitation) or land use, as well as potential microbial sources such plant cover and soil microbial composition, influence plant microbiomes (14, 15). As environmental factors influence microbial community patterns at broad geographic scales, these same factors are also perceived differentially by individual plant microhabitats (16–18). Some of these environmental factors are more relevant to belowground soil or root microbiomes, while others can be more impactful to aboveground plant microbiomes (3, 19–21). One key challenge therefore is to study microbial community dynamics in the different plant compartments simultaneously on whole plants exposed to natural environmental conditions at various sites for multiple years.

Along with environmental filtering, stochastic processes such as dispersal are also significant ecological drivers of microbial community assembly. Dispersal patterns of microorganisms across spatial and temporal scales contribute to plant microbiome dynamics, whether via horizontal transmission from the environment or vertical transmission from seeds (13, 22, 23). Changes in environmental conditions (*i.e.* nutrient resource, pH, drought, soil type) can alter likelihood of priority effects and dispersal, as well as vertical transmission of microorganisms (24–26). Survival and establishment of transmitted microorganisms in a plant compartment are then dependent on a combination of assembly factors including plant microhabitat or environmental factors (12, 27). Knowledge on the effect of environmental factors on the transmission as well as survival and persistence of microorganisms in the plant organs is critical in understanding plant microbiome dynamics. Such information will also improve our insight on how microorganisms, especially pathogens, shift from roots to aboveground plant organs or vice versa.

Thus, it is important to investigate how environmental factors shape distinct microbial communities associated with different plant organs and how these abiotic and biotic factors influence transmission and establishment of microorganisms from the roots to aboveground plant compartments. *L. corniculatus* in natural populations that are continuously exposed to natural perturbations and microbial sources is a good plant model system to study the impact of environmental factors on the structure of plant microbial communities. *L. corniculatus,* a legume species that is ubiquitous in European grasslands, can withstand a broad range of natural environments. *L. corniculatus* is a perennial flowering plant that can grow on a wide range of soil conditions, can adapt to different climates, can persist upon grazing and mowing, and hosts various insects and bees (28–31). *L. corniculatus* forms symbiotic relationships with nitrogen-fixing bacteria such as *Rhizobium japonicum*, *Mesorhizobium loti*, and *Rhizobium meliloti*, as well as arbuscular mycorrhiza fungi, which improve plant adaptation in adverse habitats (28, 29, 32, 33). *L. corniculatus* also hosts non-rhizobial endophytes that promote plant growth (34, 35). In our previous study on *L. corniculatus*, we showed the spatial and temporal variation of plant-associated microbial communities at multiple scales - the diversity and community composition vary across sampling sites and years, as well as in different plant compartments (36). We demonstrated the transmission of microorganisms between the plant organs and from other outside microbial sources which contributed to distinct yet overlapping microbial communities in the roots, shoots, flowers, and seeds (36). Although it was established that the plant organs are the primary driver of microbial community structure, it is not clear how abiotic and biotic factors as well as microbial sources from the local environment can also influence diversity and composition of the plant-associated microbial communities. Therefore, *L. corniculatus* in natural environments is an ideal system to study the roles of abiotic and biotic factors on the ecological processes that structure and maintain distinct microbial communities in different plant compartments.

In this study, we investigated the effect of environmental factors on plant-associated microbial communities. We examined the role of various abiotic and biotic factors in shaping the distinct yet overlapping microbial communities associated with roots, shoots, flowers, and seeds of *L. corniculatus* in natural populations. Specifically, we aimed (i) to establish that plant microbial community variation occur at multiple spatial and temporal scales - from sampling site and year to the level of plant organs; (ii) to show that plant microbial communities are shaped by a subset of environmental factors that are distinct with each plant organ; (iii) to identify plant organ microbe biomarkers and determine if these microorganisms differently respond to environmental factors at different plant compartments; (iv) to demonstrate that environmental factors influence transmission and establishment of microorganisms between plant organs; and finally (v) to utilize structural equation modeling to synthesize differential effects of environmental factors that contribute to structuring distinct microbial communities in plant organs. To address these objectives, we performed amplicon sequencing of microbial 16S rRNA, ITS2, and 18S rRNA genes targeting endophytic communities in roots, shoots, flowers, and seeds of *L. corniculatus* collected from seven grassland sites in the Swabian Alps, Germany for four years. Results aimed to establish an overarching understanding on how differential environmental filtering shapes the microbial communities associated with the different plant compartments.

## METHODS

### Collection and processing of *L. corniculatus*

To study the effect of local environmental factors on plant microbiomes in natural environment, we collected a total of 168 *L. corniculatus* plants and 28 soil samples from seven wild populations in the region of Swabian Alps, Germany for four years (Fig. 1a). Every year in the summer (August/September 2018-2021), we randomly sampled six plants that are flowering and fruit-producing from each of all seven sites. We took soil samples from the spot where we sampled the plants, and then pooled together for each site. To study the endophytic communities associated with the roots, shoots, flowers, and seeds, we surface-sterilized the plant organs sequentially with sterile water, epiphyte wash (1X TE + 0.1% Triton X-100), 80% ethanol, bleach (2% NaOCl), and sterile water. We stored the sterilized samples at −20 ^0^C until we process them for DNA extraction.

**Figure 1.**
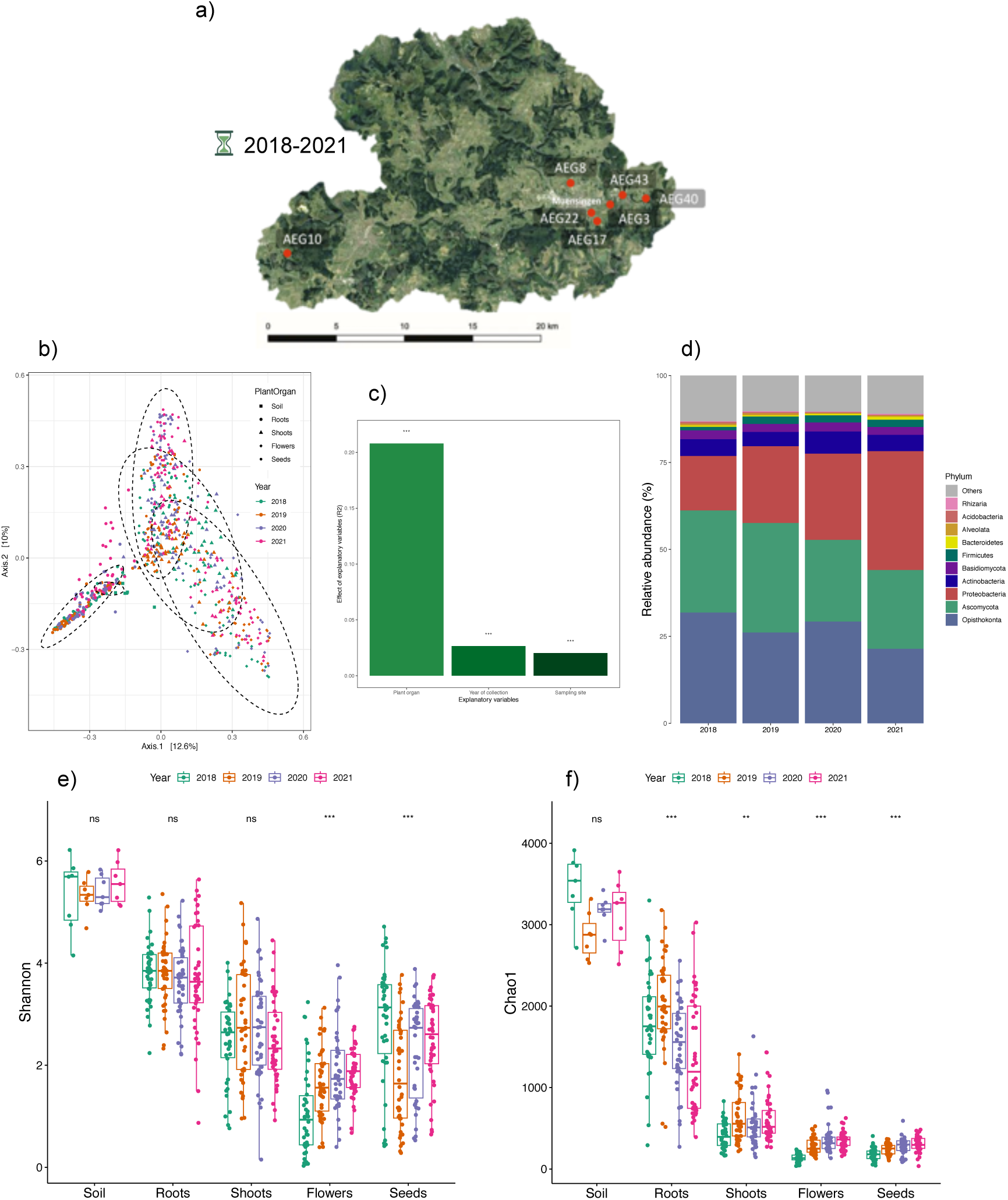
Spatial and temporal variation in *Lotus corniculatus* microbiomes. (a) Map of seven grassland sites in the Swabian Alps where plant and soil samples were collected from 2018-2021. For diversity and composition analysis, we used merged data from bacterial 16S rRNA, fungal ITS2, and eukaryotic 18S rRNA OTU tables: (b) Principal coordinate analysis based on Bray-Curtis dissimilarities between soil and plant organ microbiomes. (c) R2 statistics from PERMANOVA show the percentage of variance that can be explained by factors such as plant organ, year of collection, and sampling sites. (d) Relative abundance of ten most abundant phyla per year of collection in plants collected from seven grassland sites for four years. (e) Shannon and (f) Chao1 diversity of soil and plant organ microbiomes were compared between sampling years using Kruskal-Wallis significance test.

### DNA extraction, library preparation, and amplicon sequencing

We first homogenized frozen roots, shoots, flowers, seeds, and soil samples in Precellys 24 Tissue Homogenizer (Bertin Technologies) and then we extracted DNA using FastDNA^TM^ Spin Kit for Soil (MP Bio) following the manufacturer’s protocol. We performed two-step PCR amplification of the bacterial 16S rRNA V5-V7 region, fungal ITS2 region, and eukaryotic 18S rRNA V9 region using primers 799F/1192R, fITS7/ITS4, and F1422/R1797, respectively, on extracted DNA samples (Table S1) (37). We included blank samples (*i.e.* water and blank DNA extraction) as controls during library preparation. We also used blocking oligos, designed using R package “AmpStop”, to minimize amplification of mitochondrial and chloroplast 16S rRNA, ITS, and 18S rRNA from *L. corniculatus* (Table S1) (38). We pooled the amplification products in equimolar concentrations, purified by magnetic bead clean-up, and randomized in eight sequencing batches. We sequenced them on Illumina MiSeq with PhiX control using MiSeq Reagent Kit v3 (600-cycle).

### Processing of sequence data

We processed microbial 16S rRNA, ITS2, and 18S rRNA amplicon sequences using Mothur as described in Almario *et al.* (5, 39). In brief, we processed amplicon reads by using the following commands: *make.contigs* (contigs formation by pairing single-end reads), *screen.seqs* (quality filtering for 100-600 bases long paired reads with at least 5 bases overlap), *rename.seqs* (demultiplexing), *unique.seqs* and *count.seqs* (dereplication), *chimera.vsearch* and *remove.seqs* (detection and removal of chimera via VSEARCH), *classify.seqs* (classification of sequences), *cluster* (clustering of OTU at 97% sequence similarity threshold via the dgc method), *split.abund* (abundance filtering for OTUs with more than 50 reads), *classify.otu* (OTU classification), *make.shared* (creation of OTU tables), *remove.lineage* (removal of OTUs classified as chloroplast, mitochondria, *Arabidopsis*, *Embryophyceae*, unknown, and PhiX sequences), and *get.oturep* (identification OTU representative sequences based on abundance). We taxonomically classified bacterial 16S rRNA, fungal ITS2, and eukaryotic 18S rRNA sequences based on Greengenes database (13_8_99 release), UNITE database (02.02.2019 release), and PR2 database (version 4.12.0), respectively, which included the PhiX genome (40–42). We used Cutadapt to remove primer sequences in 16S rRNA and 18S rRNA data, while we used ITSx to remove non-ITS sequences in ITS2 data (43, 44).

### Environmental variables

This study is part of Biodiversity Exploratories (www.biodiversity-exploratories.de), an open research platform for biodiversity and ecosystem research in Germany (45). We collected samples for four years in the Swabian Alps from seven grassland sites, which have different land use types including unfertilized, mown pastures (AEG3, AEG8, AEG43), fertilized, mown pastures (AEG10, AEG40), and fertilized, mown meadows (AEG17, AEG22) (46–48). We obtained data on climatic variables (*i.e.* air temperature, precipitation, soil temperature, soil moisture) for the period of September 2017 to August 2021 from the Biodiversity Exploratories database (Biodiversity Exploratories Instrumentation Project (BExIS dataset ID 19007)). In this study we used climate values that include (i) annual mean temperature (average monthly air temperature, at 10 cm and 200 cm aboveground); (ii) temperature seasonality (amount of air temperature variation over a year, at 10 cm and 200 cm aboveground); (iii) annual precipitation (sum of all total monthly precipitation, in mm); (iv) precipitation seasonality (measure of the variation in monthly total precipitation over the course of the year, in mm); (v) soil annual mean temperature (average monthly soil temperature, at 5 cm and 10 cm below surface); (vi) soil temperature seasonality (amount of soil temperature variation over a year, at 5 cm and 10 cm below surface); (vii) annual soil moisture (sum of all total monthly soil moisture, at 10 cm below surface); and (viii) soil moisture seasonality (measure of the variation in monthly total soil moisture over the course of the year, at 10 cm below surface). For air temperature and soil temperature, we calculated the climate values from September of previous year to August of the year of sampling. For precipitation and soil moisture, we calculated climate values over the period of Water Year (*i.e.* October of previous year to September of the year of sampling). We also obtained vegetation records of the grassland sites from 2018 to 2021 from BExIS (BExIS dataset ID 31389, 20766) (49, 50). We used Shannon’s diversity and Richness indices to measure alpha-diversity and species richness of plant communities, respectively. To assess plant community composition, we performed principal component analysis (PCA) and then used PCA axis 1 and 2 scores as proxy for plant community composition. For soil microbial communities, we used Shannon’s diversity and Chao1 indices to measure the alpha-diversity and species richness, respectively. To assess soil microbial community composition, we performed principal coordinate analysis (PCoA) ordination of Bray-Curtis dissimilarities between soil samples and used PCoA axis 1 and 2 scores as proxy for soil microbial community composition. For diversity and community composition analysis of both soil microbial and plant communities, we used vegan and phyloseq R packages (51–53).

### Diversity and composition of *L. corniculatus* microbiomes

To calculate diversity, composition, and relative abundance profiles of microbial communities, we used R packages phyloseq, vegan, microbiome, and microeco to analyze OTU tables generated from Mothur sequence processing (51–55). For alpha-diversity analyses, we used Shannon’s diversity and Chao1 indices. To assess if alpha-diversity measures are significantly different between samples, we checked for normality of the data using Shapiro-Wilk normality tests, and then analyzed using the parametric test ANOVA (for normally distributed data) or the nonparametric Kruskal-Wallis rank sum test (for non-normal data). We conducted post-hoc analysis using Dunn’s test (for Kruskal-Wallis test) or Tukey’s HSD (for ANOVA test). For beta-diversity analyses, we performed Principal Coordinate Analysis (PCoA) ordination of Bray-Curtis dissimilarities between samples, then used PERMANOVA analysis of the Bray-Curtis distances to assess significant effect of explanatory variables (*i.e.* plant organ, year of collection, sampling site) on microbial community structures. We used merged data from bacterial 16S rRNA, fungal ITS2, and eukaryotic 18S rRNA OTU tables in all analyses.

### Environmental factors as explanatory variables of microbiome composition

To identify the main environmental factors that shape microbial community structure in *L. corniculatus* plant organs, we used the trans_env class of the microeco R package to analyze the merged bacterial 16S rRNA, fungal ITS2, and eukaryotic 18S rRNA datasets (55).

We inspected the significant differences of the environmental variables across sampling sites or years using ANOVA. We used Bray-Curtis distance-based RDA analysis (dbRDA) with feature selection (*i.e.* forward selection method) for explanatory environmental variables to each plant organ microbial community and calculated the adjusted R-squared of the RDA model that explains variation of community composition. We also calculated the significance of the RDA models that include the identified subset of environmental variables which best explain microbial community composition, and the contribution of such environmental variables to the model via ANOVA and vegan:envfit, respectively.

### Association of organ biomarker microbes with environmental variables

To identify bacteria, fungi, and eukaryotes that distinguish between plant organs, we used the trans_diff class of the microeco R package (55). We used linear discriminant analysis effect size (LEfSe) (*P* < 0.001, LDA score ≥ 4) to identify biomarker genera for *L. corniculatus* roots, shoots, flowers, and seeds (56). Then we used the trans_env class of the microeco R package to test for correlations between environmental variables and relative abundances of the identified biomarker microbes (55). We used heat map to visualize the Pearson correlation coefficients.

### Microbial community networks with environmental factors

To examine interactions between plant organ microbiomes and environmental factors, we performed network analysis based on correlations of OTU abundances and environmental variables. Using the trans_network class of the microeco R package, we calculated Pearson correlation coefficients of environmental variables and OTUs filtered with abundance threshold (relative abundance ≥ 0.0001) (55). We used the computed correlation coefficients (*P* < 0.0001) to build and analyse microbial community networks in Cytoscape_v3.10.1 (57). We also used the trans_env class of microeco R package to calculate correlations and to build heatmaps of bacteria, fungal, and eukaryotic genera that are highly correlated with environmental variables (55).

### Multigroup structural equation modelling

We used multigroup structural equation modelling (SEM) to understand the direct and indirect relationships of environmental factors with *L. corniculatus* microbiomes. We used a generalized linear model with gaussian distribution to investigate abiotic and biotic environmental variables as predictors of the response variable, plant microbiome composition. We grouped environmental variables into categories, including precipitation, air temperature, land use intensity, soil moisture, soil temperature, plant cover, and soil microbial diversity, and then we employed model selection to identify representative variables for multigroup SEM. For model selection, we first identified multicolinearity between variables by calculating Spearman correlation coefficients and variance inflation factors (VIF), then we excluded variables flagged as highly correlated (absolute correlation > 0.8) and with high VIF values (VIF > 5). The best-fitting model with all the representative variables was identified using stepwise selection in both forward and backward directions based on Akaike Information Criterion (AIC). To calculate standardized path coefficients, we fitted the selected GLM model in the R package piecewiseSEM with plant organs as a grouping variable and scale as type of variable standardization (58). We did all analyses in R version 4.2.1 (51).

## RESULTS

### Spatial and temporal variation in microbial communities associated with *L. corniculatus* roots, shoots, flowers, and seeds

Plant-associated microbial communities are continuously exposed to dispersal processes and filtering by local environmental conditions and host plant. These combinations of stochastic and deterministic processes occur at varying temporal and spatial scales - acting at the levels of plant compartments, above and belowground, or geographical distances, as well as at the levels of diurnal cycles to seasons. In *L. corniculatus*, plant organs are the main source of variation in the associated microbial communities (R^2^ = 0.20800, *P* < 0.001; Fig. 1bc, Fig. S1a). In addition to plant organs, sampling sites and year of collection also contributed to composition variation in bacterial, fungal, and eukaryotic communities associated with *L. corniculatus* (Fig. 1bc, Fig. S1a). The effect of year of sample collection in explaining microbial community composition variation (R^2^ = 0.02647, *P* < 0.001) is higher than sampling site (R^2^ = 0.02022, *P* < 0.001).

Significant differences in microbial community diversity are attributed to year of sample collection but not sampling site. The Shannon diversity (*P* < 0.05) of microbial communities associated with the aboveground flower and seed microbial communities vary between collection years but not in the roots and shoots (Fig. 1e). On the other hand, the diversity of the microbial communities are stable across the different sites (Fig. S1c). Meanwhile, year of sample collection contributed to significant differences in microbial community richness in all organs, and sampling site only contributed to significant differences in richness in shoot microbial communities. Chao1 richness index (*P* < 0.05) of microbial communities associated with both above- and below-ground plant organs significantly differ between collection years, and between sampling sites for shoot microbial communities (Fig. 1f, Fig. S1d).

The relative abundance profiles of the top ten bacterial, fungal, and eukaryotic phyla in the plant organs also vary between year of collection and sampling sites (Fig. 1d, Fig. S1b, Fig. S2). Therefore, the composition, diversity, and richness of microbial communities associated with *L. corniculatus* roots, shoots, flowers, and seeds are significantly affected by either spatial and temporal factors. While microbial community heterogeneity is observed between plant compartments, and especially more discernible between above-versus below-ground plant organs, observed differences in plant microbial community structures between sampling sites or years suggest that abiotic and biotic factors are acting on microbial community assembly processes at different levels of spatial and temporal scales.

### Microbial community composition in *L. corniculatus* are shaped by a subset of environmental factors that are distinct with each plant organ

To delve deeper into the processes that contribute to the observed spatial and temporal variation in *L. corniculatus*-associated microbial communities, we examined a set of environmental variables documented from seven sites (AEG3, AEG8, AEG10, AEG17, AEG22, AEG40, AEG43) of the Biodiversity Exploratories in the Swabian Alps during four years of sampling (Summers of 2018-2021) (Fig. 1a, Fig. S3-4). First, biotic and abiotic factors in the soil such as soil moisture and temperature, as well as soil microbial community composition, diversity, and richness were both taken into account (Fig. S3a-c, Fig. S4a-c). For aboveground biotic and abiotic factors, air temperature and precipitation, as well as plant community composition, diversity, and richness were analyzed (Fig. S3d-f, Fig. S4d-f). Finally, the land use intensity (LUI) in the sampling sites, including the LUI index components grazing, mowing, and fertilization, were examined (Fig. S3g, Fig. S4g). These environmental variables significantly differ between either sampling sites or years. Soil climate variables such as annual moisture, moisture seasonality, annual mean temperature, and temperature seasonality, significantly differ both among sampling sites and years. Likewise, the soil microbial community composition, diversity, and richness are also significantly different between sampling sites and years. Aboveground climate factors such as annual precipitation and annual mean temperature also vary between sampling sites and years, and climate indices such as precipitation and temperature seasonality that do not vary among sampling sites are significantly different throughout sampling years. Similarly, the composition, richness, and diversity of vegetation cover in the grassland sites vary among the seven plots, but only the plant community composition and richness significantly differ among the sampling years. The LUI of the sampling sites ranges from unfertilized and fertilized mown pastures to fertilized mown meadows. LUI and index components (*i.e.* fertilization, mowing, grazing), are significantly different between sampling sites, but not between years (except mowing).

Since each plant organ may perceive a set of environmental stimuli different from other compartments, the environmental factors that are most important in shaping the plant-associated microbial communities are most likely different across roots, shoots, flowers, and seeds. To determine the main environmental factors that are important in shaping microbial community composition in each plant organ, we applied Bray-Curtis distance-based RDA analysis (dbRDA) for explanatory environmental variables separately for each organ microbial communities. Feature selection based on forward selection method was used to identify a subset of environmental variables included in the RDA model that best explains the microbial community composition. The significance of the model and the contribution of the environmental variables to the model were also calculated (*i.e.* via ANOVA and vegan:envfit, respectively).

In general, microbial communities associated with *L. corniculatus* are shaped by all environmental factors with varying importance for each plant organ. For all plant organs, significant models (*P* < 0.05) include a subset of environmental factors that explain up to 19% variation in microbial community composition. A combination of biotic and abiotic factors explains 15% of variation in root microbial community composition (Fig. 2a). The composition of bacterial, fungal, and eukaryotic communities are mainly shaped by microbiome composition and temperature seasonality in the soil as well as richness of vegetation cover (Table S2). Meanwhile the microbial community composition in shoots are primarily driven by abiotic factors both in soil and aboveground, such as temperature and precipitation seasonality (Fig. 2b, Table S2). The set of environmental factors included in the significant model accounts for 13% of the variation in shoot microbial communities. From the model comprising a combination of factors that explains 14% of microbial community composition variation, the most important environmental drivers in shaping microbial community composition in flowers are biotic and abiotic components in the soil - microbial community composition, moisture, and temperature (Fig. 2c, Table S2). Among the set of environmental variables that account for 19% of variation, seed-associated microbial community composition are best explained by abiotic factors, such as temperature seasonality in the soil and aboveground (Fig. 2d, Table S2). Only the aboveground microbial communities are significantly affected by soil microbiome richness and diversity, as well as annual precipitation and mean temperature (Fig. 2). Plant organs are shaped by different land use indicators - root- and flower-associated microbial communities are significantly affected by fertilization, while shoot and flower microbiomes are significantly shaped by mowing and grazing, respectively (Fig. 2). Therefore, variation in microbial communities associated with different plant organs are shaped by a set of environmental perturbations that are distinct to each plant compartment.

**Figure 2.**
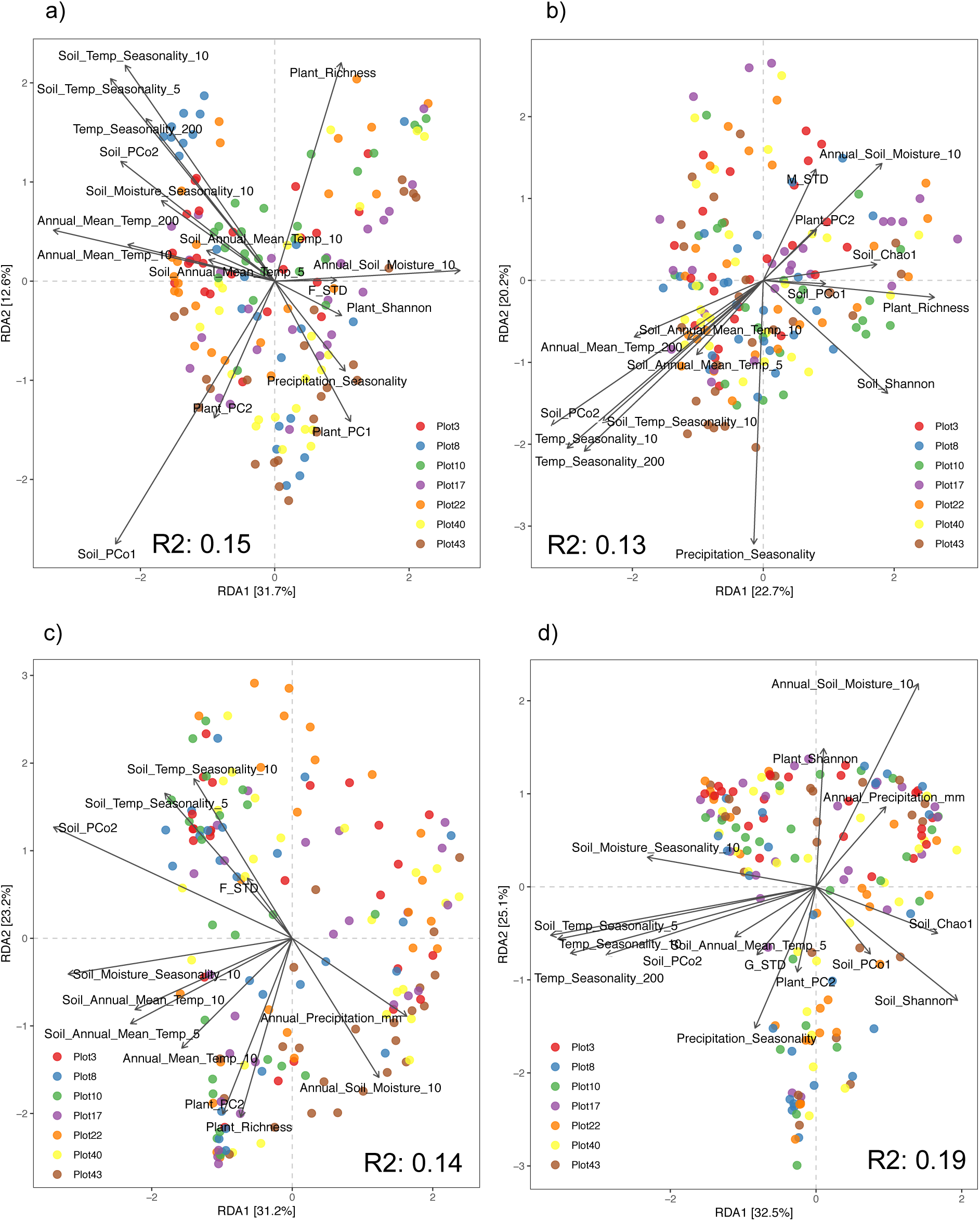
*Lotus corniculatus* microbiomes are shaped by a subset of environmental variables that are different with each plant organ. Bray-Curtis distance-based RDA analysis (dbRDA) of (a) root, (b) shoot, (c) flower, and (d) seed microbiomes, with vectors representing environmental variables which were feature-selected based on forward selection method. Variance (adjusted R^2^) of microbial communities explained by the feature-selected environmental variables included in the significant RDA models (*P* < 0.05) are indicated.

### Organ biomarker microbes selectively respond to environmental factors in different *L. corniculatus* organs

Plant-associated microbial communities below- and aboveground are exposed to similar local environmental conditions but are perceived at different scales, and these perturbations, in addition to other plant host selection and stochastic processes, consequently lead to the distinct but overlapping community structures of microbial communities associated with the roots, shoots, flowers, and seeds (Fig. 1). To probe if environmental factors are perceived by microorganisms differently in different plant compartments, we assessed if the abundance of organ biomarker microbes are correlated with environmental factors. Biomarker microbes, which are differentially abundant microbes that account for the differences between plant organ microbial communities, were identified via LEfSe analysis (Fig. 3ace). The relationship between the abundance in each plant organ of these biomarker microbes and environmental variables were calculated using Pearson correlation.

In the roots, the abundance of root biomarker bacteria *Mesorhizobium* and *Cryptosporangium* significantly correlated with environmental variables, but showed no significant correlation in other plant organs (Fig. 3b, Fig. S6). Most of the environmental variables important in shaping root microbial community composition, such as temperature seasonality in the soil and aboveground air, soil moisture seasonality, and soil microbial composition, are negatively correlated with the abundance of *Mesorhizobium* in the roots (Fig. 2a). *Cryptosporangium* abundance are significantly correlated with soil moisture and plant diversity, which are variables that also significantly shape root microbial communities (Fig. 2a). In the shoots, shoot biomarker fungi *Setosphaeria* abundance is positively correlated with soil microbial community composition, while in the roots their abundance is negatively correlated with soil microbial community richness and diversity (Fig. 3d, Fig. S7). These biotic factors (*i.e.* soil microbes) are relevant factors in shaping shoot microbial community composition (Fig. 2b). Abundance of flower biomarker microbes *Wolbachia* and *Botryotinia* are significantly correlated with environmental variables in the flowers but not in other plant organs (Fig. 3ac, Fig. S6-7). *Wolbachia* abundance is positively correlated with microbial community composition and moisture seasonality in the soil, temperature seasonality both in the soil and aboveground, and precipitation seasonality - all important variables influencing microbial communities of all plant organs (Fig. 2). *Botryotinia* abundance also significantly correlated with microbial community composition and temperature seasonality in soil as well as temperature and precipitation aboveground.

**Figure 3.**
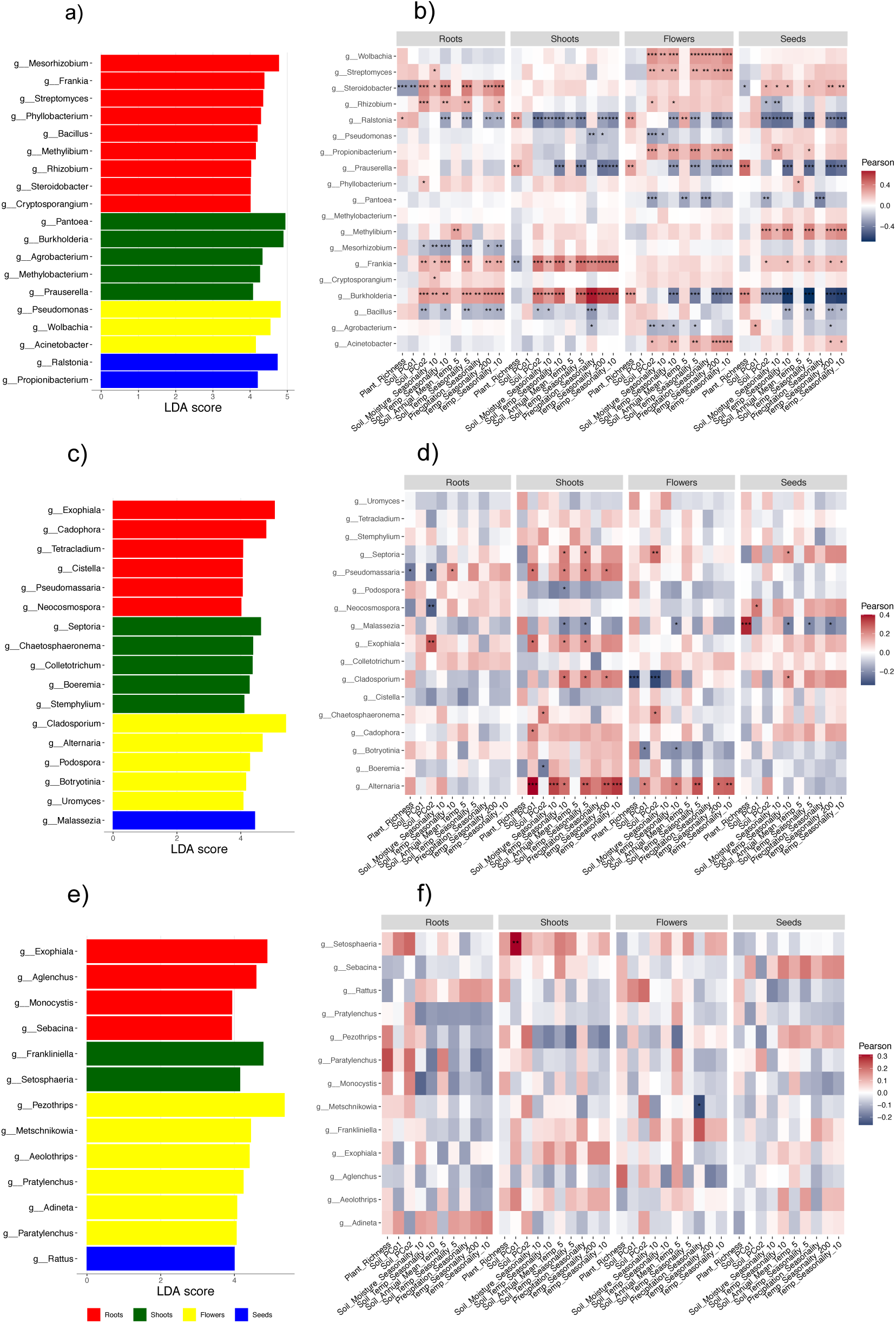
*Lotus corniculatus* organ biomarker microbes respond differently to environmental variables in different plant organs. LEfSe analysis to identify differentially abundant (a) bacterial, (c) fungal, and (e) eukaryotic biomarkers (*P* < 0.001, LDA score ≥ 4) of the plant organ microbiomes. Heatmap of Pearson correlations between selected environmental variables and relative abundances of the identified (b) bacterial, (d) fungal, and (f) eukaryotic organ biomarker microbes in roots, shoots, flowers, and seeds.

The abundance of aboveground biomarker microbes for the shoot (*i.e. Prauserella, Pantoea, Agrobacterium, Septoria, Chaetosphaeronema*), flower (*i.e Pseudomonas, Cladosporium, Alternaria*), and seed (*i.e. Propionibacterium, Malassezia*) microbial communities are significantly correlated with environmental factors in the aboveground plant organs but not in the roots (Fig. 3, Fig. S6-8). Meanwhile there are also biomarker microbes, such as the shoot biomarker *Burkholderia* and seed biomarker *Ralstonia,* in which their abundance in all plant compartments are significantly associated with environmental factors (Fig. 3b, Fig. S6). Some microorganisms that are important members of the plant organ microbial communities respond to environmental perturbations only in the corresponding plant organ but not in other compartments, therefore highlighting the role of a set of environmental variables that are important in shaping plant organ microbial communities but otherwise have weak effect in other plant compartments. Many aboveground biomarker microorganisms are also receptive to environmental factors only in aboveground compartments and not in the roots. On the other hand, some biomarker microorganisms perceive and are affected by such environmental variables anywhere in the plants.

### Organ-specific filtering by environmental factors influence transmission and establishment of microorganisms in different *L. corniculatus* organs

Distinct *L. corniculatus* organ microbiomes are overlapping and interconnected via transmission of microorganisms between plant compartments and from outside environment (36). The microbial communities in roots, shoots, flowers, and seeds are shaped by a set of environmental factors that are distinct to each plant compartment. To examine how environmental factors can influence transmission of microorganisms within plants and how these factors contribute to assembly of these microbial communities to become organ-specific, we analyzed how local biotic and abiotic factors shape community structure of plant organs, specifically looking into how these factors impact microbial interactions in the different plant organs. To analyze interactions between environmental factors and plant organ microbiomes, we performed network analysis based on Pearson correlation coefficients of OTU abundances and environmental variables and analyzed co-occurrence patterns between microbial taxa and abiotic and biotic factors (Fig. 4). After applying abundance threshold (relative abundance ≥ 0.0001) and criteria for correlation (*P* < 0.0001), the complexity of root microbial network is highest with 708 nodes and 14,977 edges, followed by shoot microbial network with 338 nodes and 4,014 edges, seed microbial network with 159 nodes and 1,358 edges, and finally flower microbial network with the least number of nodes (119) and edges (429) (Table S3). The root microbial network has more positive correlations (73% of total significant correlations) than negative correlations (27%) with environmental variables, while the shoot microbial network is evenly positively and negatively correlated (51% and 49%, respectively), and both the flower and seed microbial networks have less positive correlations (20% and 26%, respectively) than negative correlations (80% and 74%, respectively) with environmental variables.

**Figure 4.**
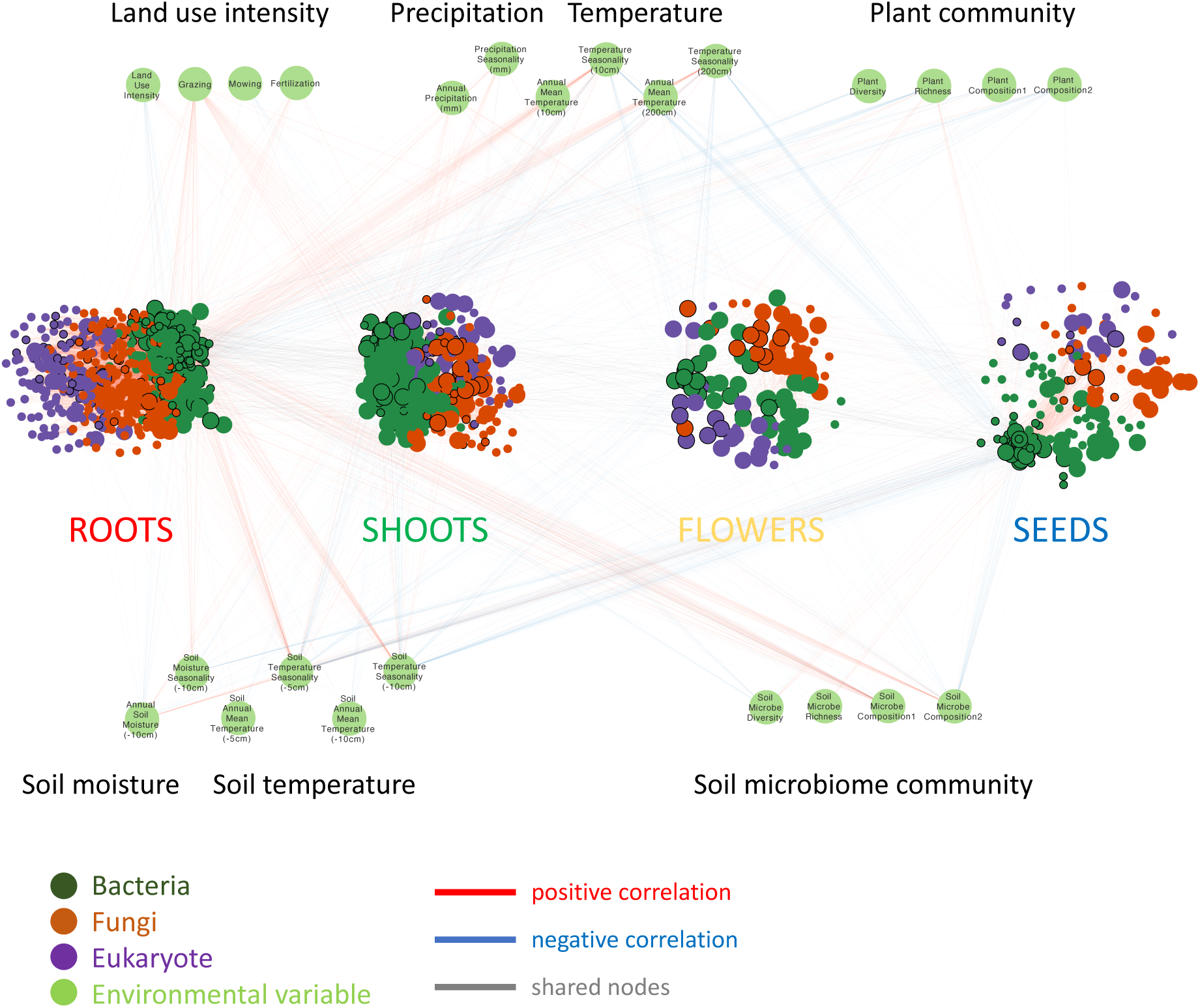
Environmental factors influence microbial interactions in *Lotus corniculatus* organ microbiomes. Root, shoot, flower, and seed networks based on Pearson correlations (*P* < 0.0001) of OTU abundances and environmental variables. Bigger-sized nodes are shared between adjacent plant compartments and nodes with wider border are significantly correlated with environmental variables.

The environmental factors that impact the structure of microbial community networks in the plants differentially affect the plant compartments (Fig. 4). Thirty percent of the OTUs in the plant microbiome network are significantly correlated with environmental variables (*i.e*. specifically 29%, 29%, 31%, and 35% of the OTUs in the root, shoot, flower, and seed microbiome networks, respectively). Among the soil climate variables, soil temperature seasonality has the highest number of significant correlations. Most of the correlations between soil temperature seasonality and microbes in the roots are positive, while in the aboveground plant parts the correlations are negative. Soil microbiome composition also showed high number of correlations and is mostly positively correlated with root microbes and negatively correlated with microbes in aboveground organs. Among the climate variables aboveground, temperature seasonality has the most number of significant correlations. In the roots and shoots, temperature seasonality is primarily positively correlated with the microbes, in contrast it is negatively correlated with microbes in the flowers and seeds. Plant community richness is also highly correlated with the microbes in the plant, and while it is mostly negatively correlated with the root microbes, in the aboveground plant parts it is primarily positively correlated. The relationship between LUI and microbe abundances differs across different plant organs - it is negatively correlated in the roots, positively correlated in the shoots, and no correlation in flowers and seeds. Among the LUI components, grazing has the highest number of correlations with the plant microbes, and is positively correlated with the microbes in all plant organs. Overall, soil temperature seasonality, soil microbiome composition, temperature seasonality, plant community richness, and grazing are the most relevant environmental variables in the microbial networks, and such environmental variables often differ in relationships with microbes in the roots compared with aboveground microbes in shoots, flowers, and seeds.

Throughout continual transmission of microorganisms across plant compartments, the microorganisms that eventually survive and persist in each organ depend on a combination of host and environment conditions, including abiotic and biotic factors. The contrast in relationship between environmental factors and microbe abundances in a particular organ compared with other plant organs possibly played essential role in shaping the distinct plant organ communities. In the root network, 19% of the OTUs are potentially transmitted to the shoot network, while 26% of the shoot network OTUs are transmitted to the flower network, and 51% of the flower network OTUs goes to the seed network (Fig. 4). Among the transmitted microbes to the shoot network, 34% are significantly correlated with environmental variables, while 19% and 36% of the microbes that goes to the flower and seed networks, respectively, have significant interactions with environmental variables. For instance, *Burkholderia* and *Sulfuritalea*, which are microorganisms that are present in all organs and thus are potentially transmitted across organs, are highly correlated with environmental variables (Fig. S9). *Burkholderia* and *Sulfuritalea* abundances are positively correlated with soil temperature seasonality, soil microbiome composition, and temperature seasonality in the roots and shoots, and shifted to negative relationship in flowers and seeds. Meanwhile their abundances are positively correlated with plant community richness in the flowers and seeds. This is consistent with the observed patterns that the association between microbial abundances and environmental factors shifts when microorganisms are transmitted from roots to aboveground plant compartments.

### Structural equation modeling infers direct and indirect environmental effects that shape microbial community composition in *L. corniculatus* organs

The multigroup SEM, with plant organ as grouping variable, aims to estimate direct and indirect effects of environmental factors on microbial community composition in roots, shoots, flowers, and seeds (Fig. 5). Through SEM we assessed the direct influence of abiotic (precipitation, air and soil temperature, soil moisture, and LUI) and biotic (community composition of plant cover and soil microbiome) environmental factors on plant microbial community composition, as well as indirect influence of the abiotic factors via their effect on such biotic factors (Fig. 5a). We built a SEM model (Fisher’s C = 2.032, *P* = 0.362, Fig. 5b) that estimated the direct effect of the abiotic variables - annual precipitation, temperature seasonality at 200 cm aboveground, LUI, annual soil moisture at 10 cm below surface, and soil annual mean temperature at 10 cm below surface - on plant microbial community composition (PCoA axis 2 scores). The indirect effect of these abiotic variables through their influence on the biotic variables - plant cover composition (PCA axis 2 scores) and soil microbiome community composition (PCoA axis 2 scores) - was also evaluated.

**Figure 5.**
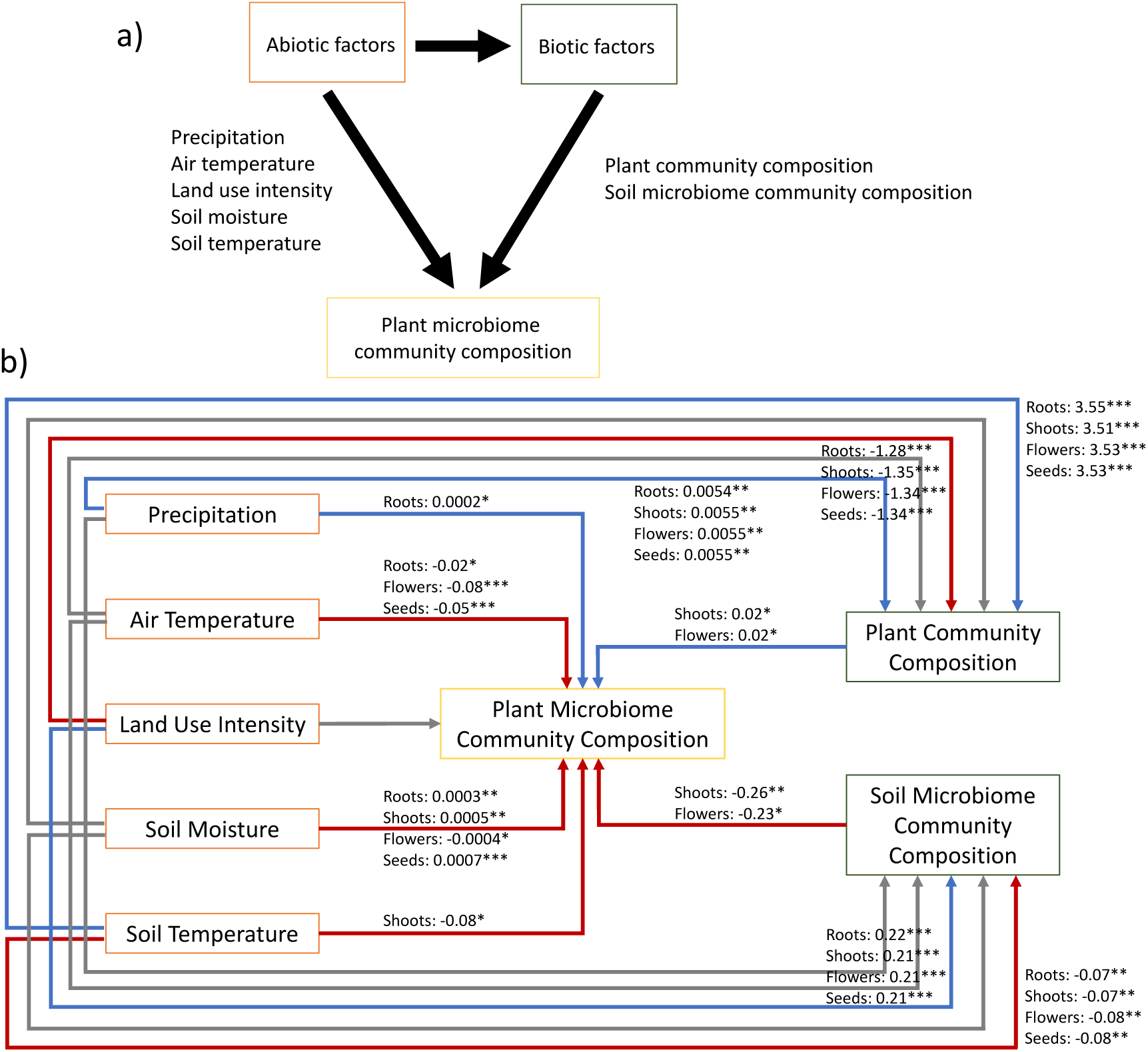
Multigroup SEM shows direct and indirect effects of environmental factors on *L. corniculatus* organs. (a) Causal hypothesis for the multigroup SEM which proposes the direct and indirect (via biotic factors) effects of abiotic environmental factors on plant microbiome community composition. b) Multigroup SEM, with plant organs as grouping variable, show significant path coefficients to estimate the effects of precipitation (annual precipitation), air temperature (temperature seasonality at 200 cm aboveground), land use intensity (LUI), soil moisture (annual soil moisture at 10 cm below surface), soil temperature (soil annual mean temperature at 10 cm below surface), plant cover composition (PCA axis 2 scores), and soil microbiome community composition (PCoA axis 2 scores) on plant microbiome community composition (PCoA axis 2 scores). Blue arrows indicate positive and red arrows indicate negative path coefficients. Gray arrows indicate nonsignificant path coefficients but are included in the significant model. Only significant path coefficients are shown: ***, *P* < 0.001; **, *P* < 0.01; *, *P* < 0.05.

We found that precipitation and plant community composition directly increase community composition of plant microbiomes, while temperature (air and soil) and soil microbiome community composition negatively affect plant microbiome community composition. Soil moisture positively affects microbiome community composition in roots, shoots, and seeds, but negatively influences flower microbiomes. LUI has no direct effect on plant microbiomes, but it can indirectly influence them via its negative and positive relationship with plant cover and soil microbiomes, respectively. Thus, through multigroup SEM we establish that environmental factors differentially affect the different plant organs and possibly dispersal decisions of microorganisms. The calculated path coefficients, although significant, are low and may explain a small proportion of variation in plant microbiome community composition.

## DISCUSSION

*L. corniculatus* microbial communities are distinct yet overlapping across roots, shoots, flowers, and seeds. In addition to organ-specificity of the microbiomes, we observed spatial and temporal variations across multiple sites and years. While plant organs are the primary source of microbiome variation, this study aims to further explore other factors that account for the observed spatial and temporal variations. Specifically, we examined how the abiotic and biotic factors in the local environment throughout multiple years contribute to the structure of microbial communities associated with *L. corniculatus* populations. We observed that plant organ microbial communities are shaped by a set of environmental factors that are distinct to each plant compartment. These environmental factors influence plant microbiomes, either by directly affecting them or indirectly through their influence on other biotic factors. Dispersal of microorganisms within plant compartments and from surrounding environment, and afterwards organ-specific filtering by biotic and abiotic factors, contribute in shaping distinct and overlapping microbial communities across plant organs. Subsequent establishment and persistence of microbial communities after dispersal are dependent on their differential perception of environmental conditions in each plant organ.

In general, *L. corniculatus* microbial communities are shaped by every local abiotic and biotic factor with varying importance for each plant organ. In the roots, both abiotic and biotic factors in the soil, such as microbiome composition and temperature, primarily explain variation in microbial communities. It is established that soil microbiomes are the primary reservoir of microorganisms that are recruited towards and enter the roots (10, 19, 59). Temperature changes in the soil directly influence root microbial community composition by regulating which microbial taxa survive and dominate in the community and indirectly by shifting the community structure of the soil microbial sources and by impacting recruitment of microbes due to changes in root exudation patterns (60–62). Vegetation cover is also found to influence soil and root microbial communities mainly by affecting soil properties (63, 64). In the shoots, abiotic factors temperature and precipitation largely explain microbial community structure. Temperature, both in soil and aboveground, significantly impacts leaf microbiomes by shifting increase or decrease of beneficial and pathogenic microbial taxa (65–68). Similarly, precipitation also impacts abundance of plant pathogens, and can serve as microbial reservoir of phyllosphere microbiota (69, 70). Microbial communities in the flowers are primarily explained by both abiotic and biotic components of the soil. Soil microbes have been found to reach and colonize flower tissues (19, 71). Conditions in the soil like temperature shape soil microbiomes as well as impact recruitment and selection of microbes by the roots and thus indirectly filter microbes that eventually reach the flowers (72, 73). Generally, changes in abiotic conditions like temperature and moisture induce changes in vegetative and reproductive plant tissues and their associated microbial communities (74). Seed microbial communities are mainly affected by temperature, both in soil and aboveground. The environment including abiotic components like temperature affects the structure of seed-associated microbial communities by significant enrichment of microbial taxa leading to changes in co-occurrence patterns (75, 76). Furthermore, LUI components fertilization, mowing, or grazing differently affect *L. corniculatus* organs. Microbial communities in roots are impacted by fertilization, while shoot microbial communities are affected by mowing. Nitrogen fertilization regulates soil microbe abundance, root exudation, and microbe recruitment resulting in enrichment of microbial groups and genes involved in nitrogen cycle in the roots (77–79). Long-term mowing impacts leaf microbial community structure with variation in leaf functional traits and enrichment of microbial taxa like Actinobacteria (80). Flower microbial communities are influenced by fertilization and grazing. Fertilization can modify the chemical properties such as N and C availability in the phyllosphere, and is found to alter microbial community diversity in flowers associated with enriched microbial groups (81). Disturbances like grazing also alter flower microbial communities by creating stress and physical changes in the flower microhabitats or by introduction of microbes by grazing animals, resulting in shift in abundances of some microbial genera (81).

In the *L. corniculatus* endophytic metacommunities, transmission of microorganisms between plant organs is determined by environmental factors along with the differing organ microhabitats and microbial interactions, resulting in distinct yet overlapping microbial communities of the roots, shoots, flowers, and seeds that are linked by dispersal. Organisms evaluate their dispersal decisions based on conditions suitable for their survival and reproduction, considering ecological cues such as abiotic conditions and species interactions (82). In *L. corniculatus*, the environmental factors soil temperature seasonality, soil microbiome composition, air temperature seasonality, plant community richness, and grazing, which are found to influence the structure and microbial interactions in the organs and are different in association per organ microbial communities, possibly influence dispersal decisions of microorganisms whether to settle or leave the plant compartments. Indeed, soil microbial communities, temperature in soil and aboveground, plant communities, and grazing are important in structuring plant-associated microbial communities (19, 59, 63–68, 81, 83, 84). *Burkholderia*, a genus that includes plant-associated microbes that can either be pathogenic, neutral, or beneficial to their hosts, are microbes in *L. corniculatus* that are highly correlated with environmental variables across all plant organs and respond to such environmental variables differently depending on their organ microhabitat (85). *Burkholderia* abundance fluctuate in response to environmental factors depending on the habitat, for instance, their abundance are positively associated with temperature in soil and aboveground as well as soil microbiome composition in the roots and shoots while negatively correlated in the flowers and seeds. This organ-dependent environmental perception is also observed in the rhizobial symbiont *Mesorhizobium*, leaf pathogen *Setosphaeria*, and necrotroph *Botryotinia*, which are biomarker microbes in roots, shoots, and flowers, respectively (86–88). Such biomarker microbes are abundant and persistent in their corresponding organ microhabitats and their habitat choice is influenced by their differential response to environmental factors in different plant compartments. It is established that abiotic and biotic perturbations can impact chemical and morphological properties in plant compartments, thus can explain the differential response of microbes to environmental factors in different plant organs (89, 90). Therefore, dispersal decisions or habitat choice, and consequently structure of microbial communities, depend on the interactions of microhabitats, key microbes, and environmental factors. This also highlights the role of a set of environmental variables that are important in shaping microbial communities in a plant organ that is different compared with other plant compartments.

It has been previously shown that LUI impacts bacterial diversity in roots, leaves, and flowers, however in our study the effect of the individual LUI component grazing on *L. corniculatus* microbiome structure is more pronounced, probably due to low variation in LUI among the seven sampling sites (81, 91). Through SEM approach we detect the indirect influence of LUI on plant microbiomes via their effects on plant cover and soil microbiomes. While we examined soil and aboveground abiotic (*i.e.* temperature and moisture/precipitation) and biotic factors (*i.e.* soil microbiomes, vegetation cover), as well as LUI (*i.e.* fertilization, grazing, mowing), other environmental variables like humidity, solar radiation, or edaphic factors such as soil pH and soil C:N also influence plant microbiomes (19, 92, 93). The observed patterns on dispersal decisions or habitat choice based on organ-dependent environmental cues and microbial interactions in *L. corniculatus* microbiomes advance our insights on how beneficial microbes or pathogens survive and reproduce on specific plant microhabitat and environmental conditions, providing basis for further investigations to test how to mitigate pathogen spread or to engineer perturbation-resistant microbiomes. Our knowledge on the organ-specific response of plant microbiomes to abiotic and biotic factors will equip us with a framework to understand and manipulate plant microbiomes experiencing climate change.

## ACKNOWLEDGEMENTS

We thank Bossdorf Lab and Kemen Lab for participating in the sampling trips, especially Frank Reis and Elke Klenk for helping in collecting and processing the samples. Through a cooperation agreement with Biodiversity Exploratories (DFG Priority Program 1374) we were granted access to the Schwäbische Alb plots. We thank Ralf Lauterbach and Jörg Hailer from the Schwäbische Alb local management team of the Biodiversity Exploratories for instructions and support during the field work. We thank the managers of the Schwäbische Alb Exploratory, Kirsten Reichel-Jung and Julia Bass and all former managers for their work in maintaining the plot and project infrastructure, Victoria Grießmeier for giving support through the central office, Andreas Ostrowski for managing the central database, and Markus Fischer, Eduard Linsenmair, Dominik Hessenmöller, Daniel Prati, Ingo Schöning, François Buscot, Ernst-Detlef Schulze, Wolfgang W. Weisser and the late Elisabeth Kalko for their role in setting up the Biodiversity Exploratories project. We thank the administration of the UNESCO Biosphere Reserve Swabian Alb as well as all land owners for the excellent collaboration. Field work permits were issued by the responsible state environmental office of Baden-Württemberg.

## AUTHOR CONTRIBUTIONS

All authors conceptualized this work. K.L. collected and processed samples from the field, and performed the experiments and data analysis. K.L. wrote the manuscript with contributions from all authors. All authors read and approved the final manuscript.

## COMPETING INTERESTS

We declare no competing interest in relation to this work.

## FUNDING

This project has been funded by the DFG program SPP 2125 DECRyPT and the ERC program DeCoCt.

## Notes

### Competing Interest Statement

The authors have declared no competing interest.

