## Supplementary Figures and Tables for "Organ-specific filtering by abiotic and biotic environmental factors shapes distinct yet overlapping microbial communities across *Lotus corniculatus* roots, shoots, flowers, and seeds"

### SUPPLEMENTARY MATERIALS

### SUPPLEMENTARY FIGURES

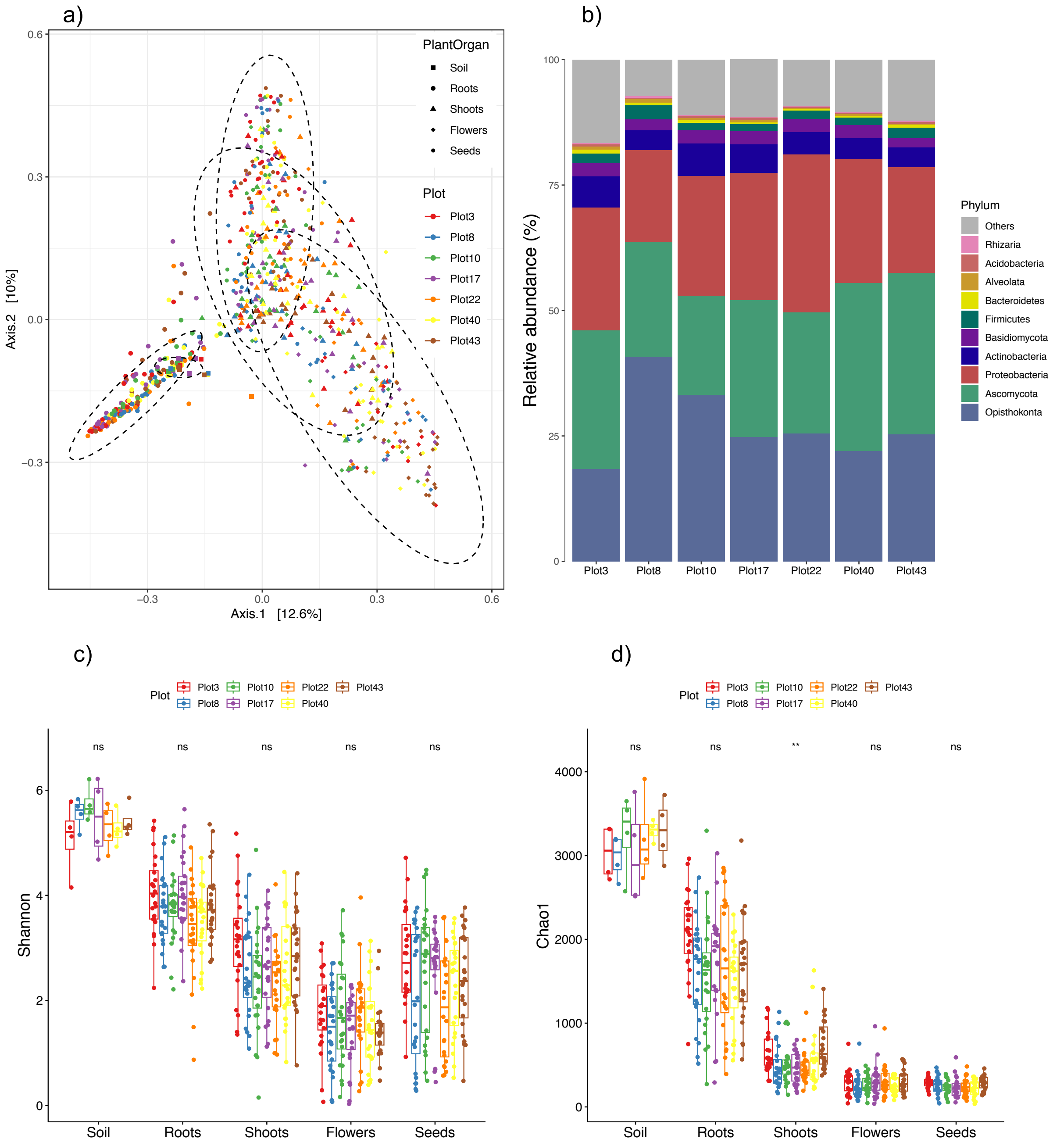


**Supplementary Figure 1.** Diversity and composition analysis based on merged data from bacterial 16S rRNA, fungal ITS2, and eukaryotic 18S rRNA OTU tables. (a) Principal coordinate analysis based on Bray-Curtis dissimilarities between soil and plant organ microbiomes. (b) Relative abundance of ten most abundant phyla per sampling site in plants collected from seven grassland sites for four years. (c) Shannon and (d) Chao1 diversity of soil and plant organ microbiomes were compared between sampling sites using Kruskal-Wallis significance test.

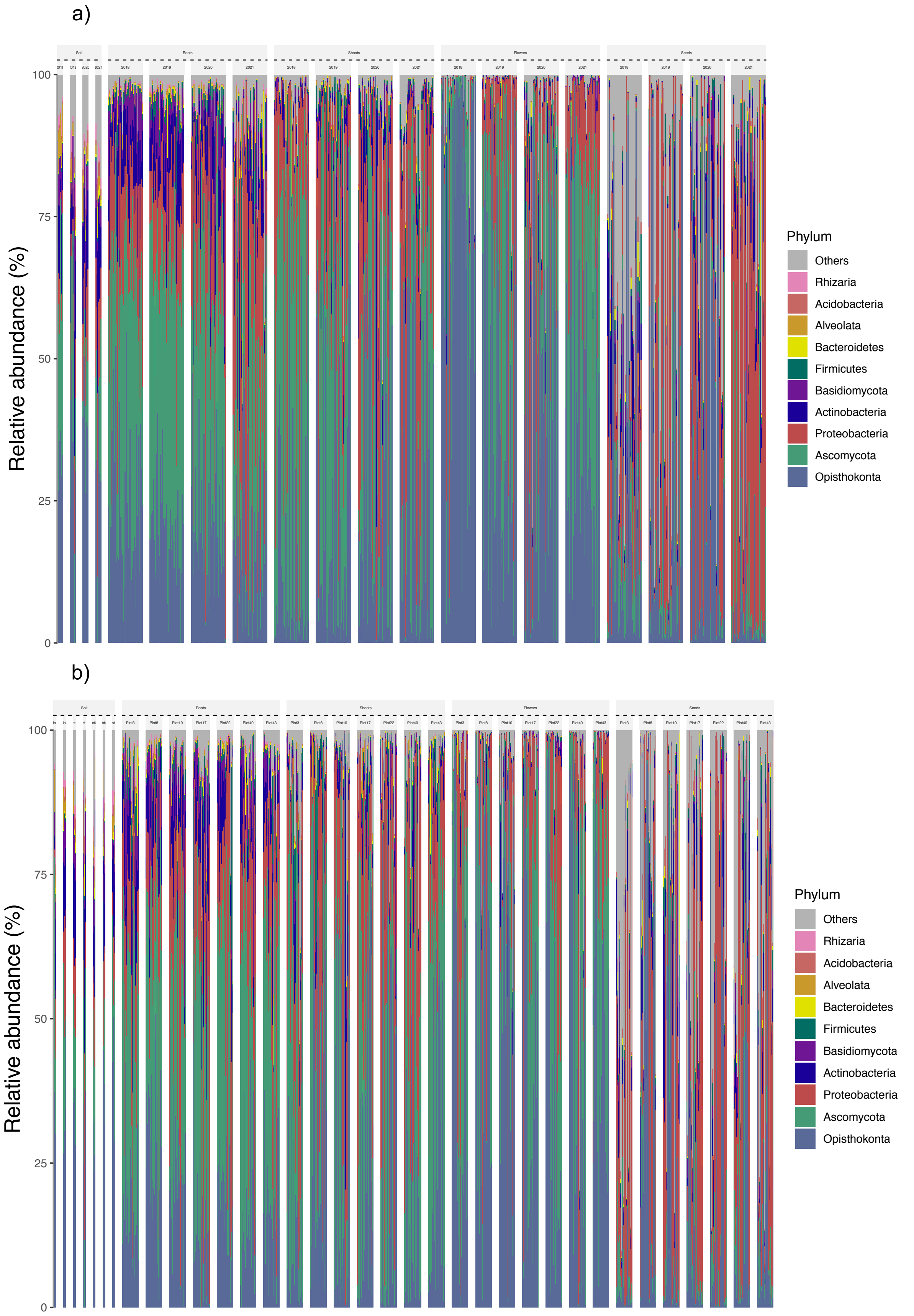


**Supplementary Figure 2.** Relative abundance of ten most abundant phyla in soil, root, shoot, flower, and seed samples collected for (a) four years from (b) seven grassland sites.

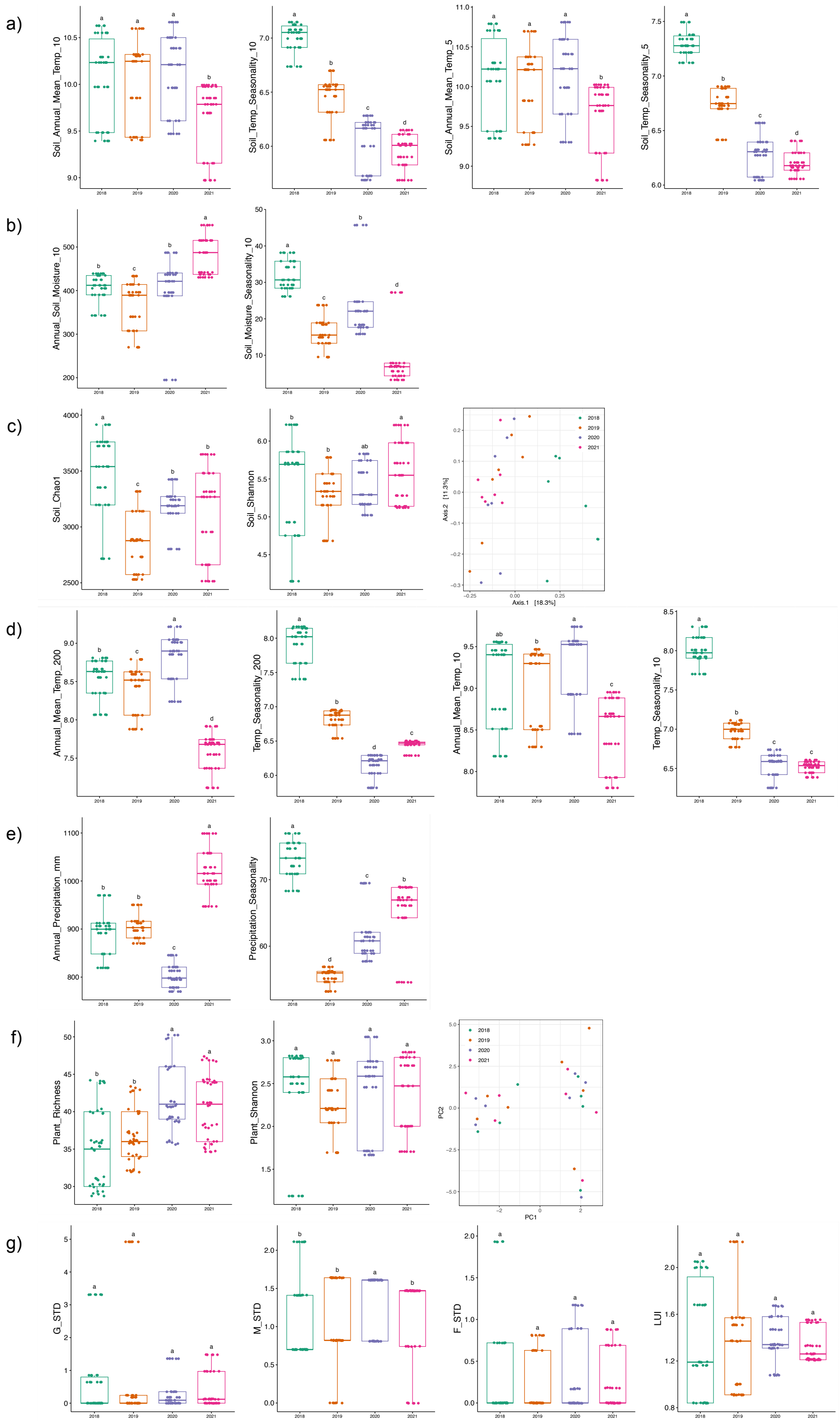


**Supplementary Figure 3.** Local environmental conditions during four years of sampling. (a) Soil temperature (soil annual mean temperature, soil temperature seasonality, at 5 cm and 10 cm below surface); (b) soil moisture (annual soil moisture, soil moisture seasonality, at 10 cm below surface); (c) soil microbiome composition (Shannon's diversity, Chao1 indices, PCoA); (d) air temperature (annual mean temperature, temperature seasonality, at 10 cm and 200 cm aboveground); (e) precipitation (annual precipitation, precipitation seasonality); (f) plant community composition (Shannon's diversity, Richness, PCA); and (g) land use intensity (fertilization, grazing, mowing) were compared between years of collection using ANOVA.

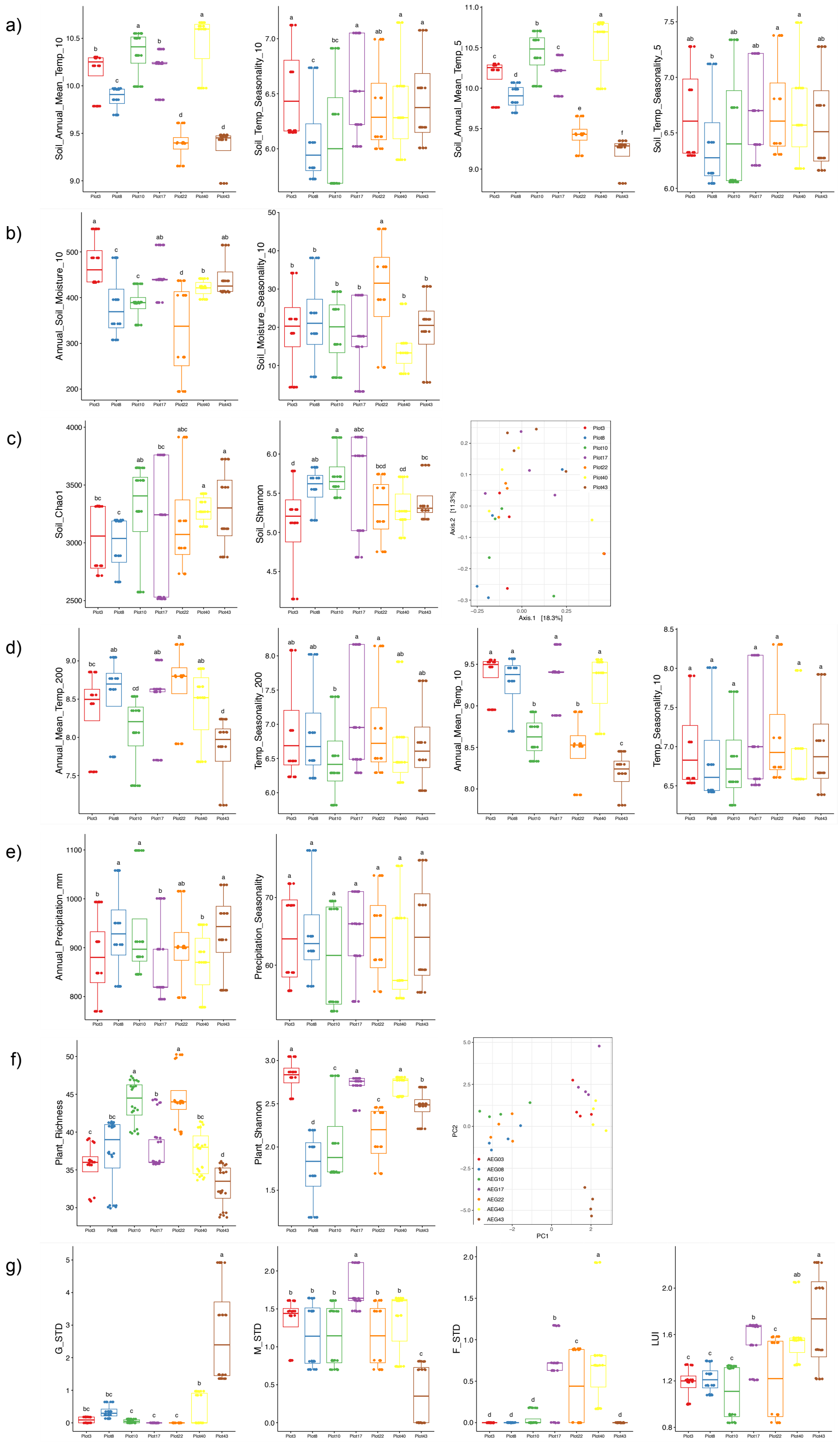

**Supplementary Figure 4.** Local environmental conditions across seven grassland sites. (a) Soil temperature (soil annual mean temperature, soil temperature seasonality, at 5 cm and 10 cm below surface); (b) soil moisture (annual soil moisture, soil moisture seasonality, at 10 cm below surface); (c) soil microbiome composition (Shannon's diversity, Chao1 indices, PCoA); (d) air temperature (annual mean temperature, temperature seasonality, at 10 cm and 200 cm aboveground); (e) precipitation (annual precipitation, precipitation seasonality); (f) plant community composition (Shannon's diversity, Richness, PCA); and (g) land use intensity (fertilization, grazing, mowing) were compared between sampling sites using ANOVA.

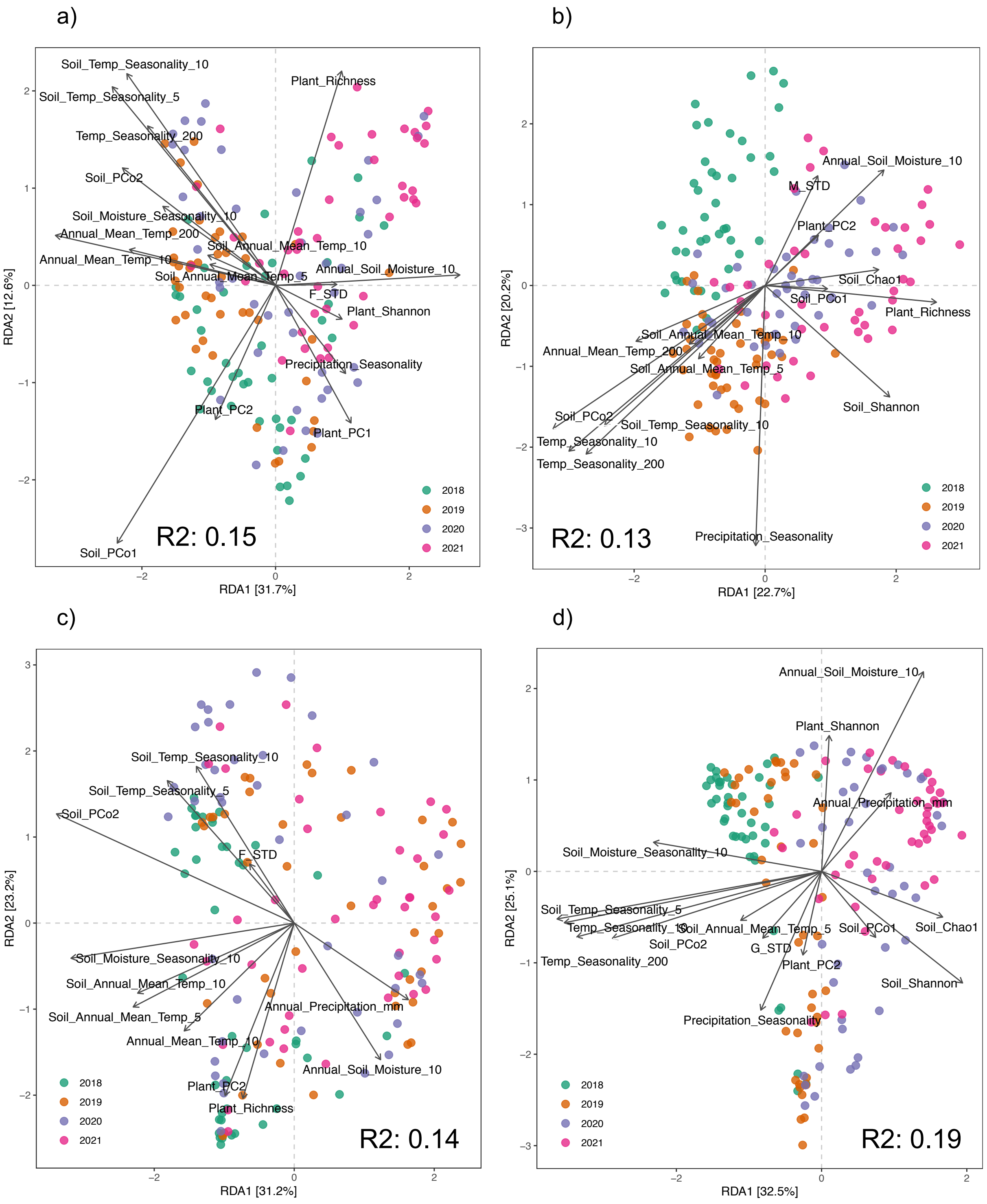


**Supplementary Figure 5.** Bray-Curtis distance-based RDA analysis (dbRDA) of (a) root, (b) shoot, (c) flower, and (d) seed microbiomes, with vectors representing environmental variables which were feature-selected based on forward selection method. Variance (adjusted $R^2$) of microbial communities explained by the feature-selected environmental variables included in the significant RDA models ($P < 0.05$) are indicated.

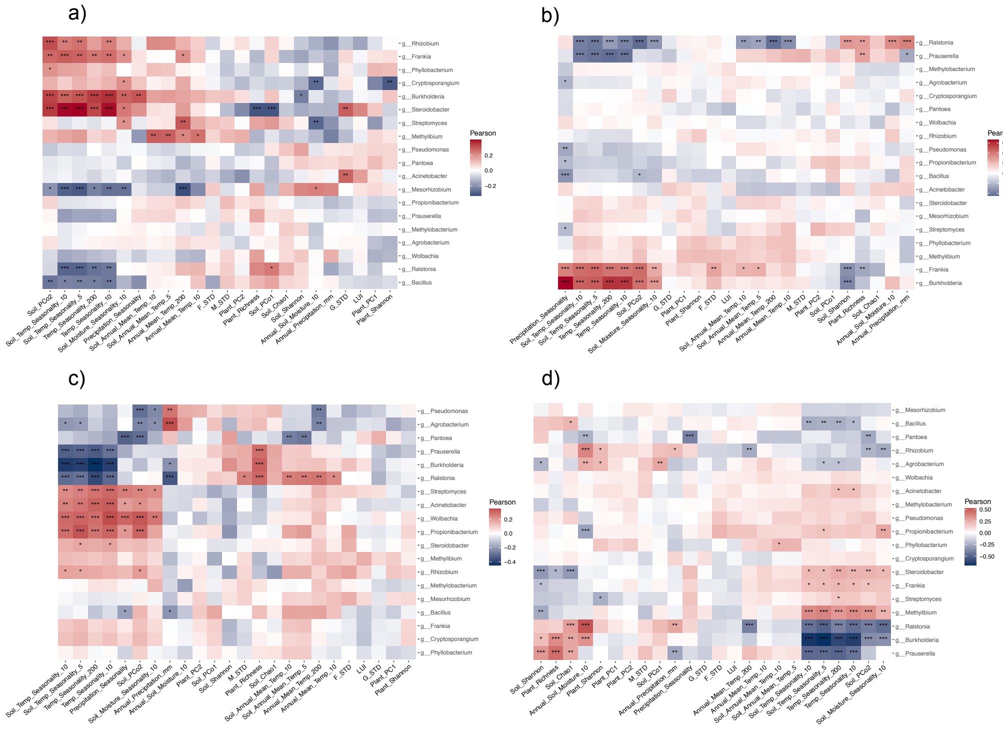

**Supplementary Figure 6.** Heatmap of Pearson correlations between all environmental variables and relative abundances of the identified bacterial organ biomarker microbes (in Fig. 3a) in (a) roots, (b) shoots, (c) flowers, and (d) seeds.

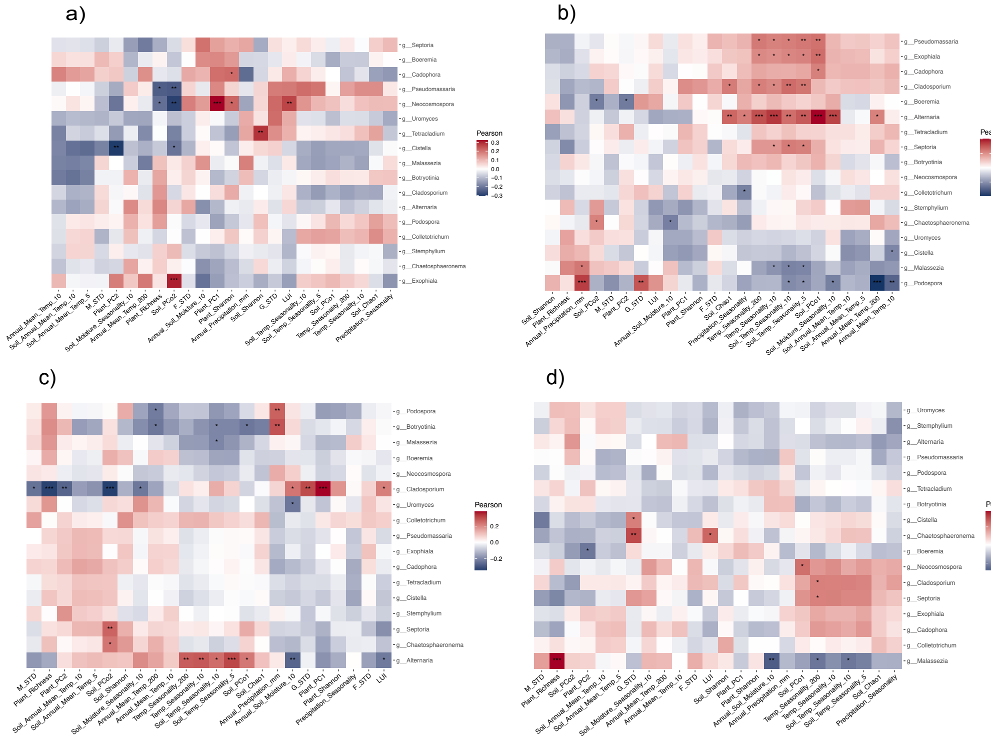

**Supplementary Figure 7.** Heatmap of Pearson correlations between all environmental variables and relative abundances of the identified fungal organ biomarker microbes (in Fig. 3c) in (a) roots, (b) shoots, (c) flowers, and (d) seeds.

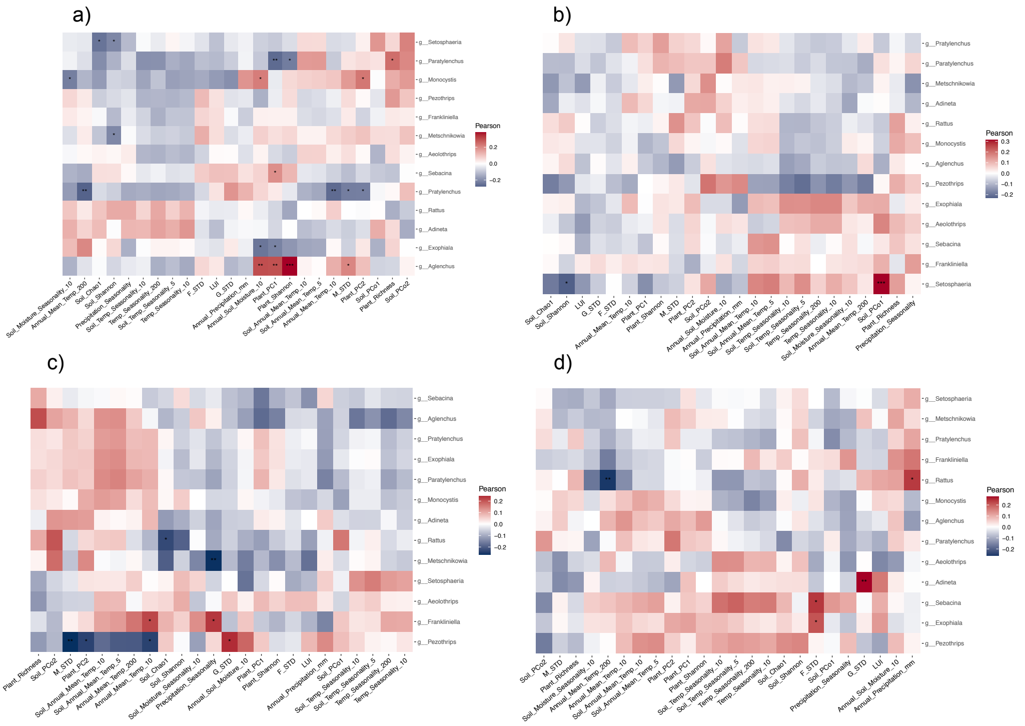

**Supplementary Figure 8.** Heatmap of Pearson correlations between all environmental variables and relative abundances of the identified eukaryotic organ biomarker microbes (in Fig. 3e) in (a) roots, (b) shoots, (c) flowers, and (d) seeds.

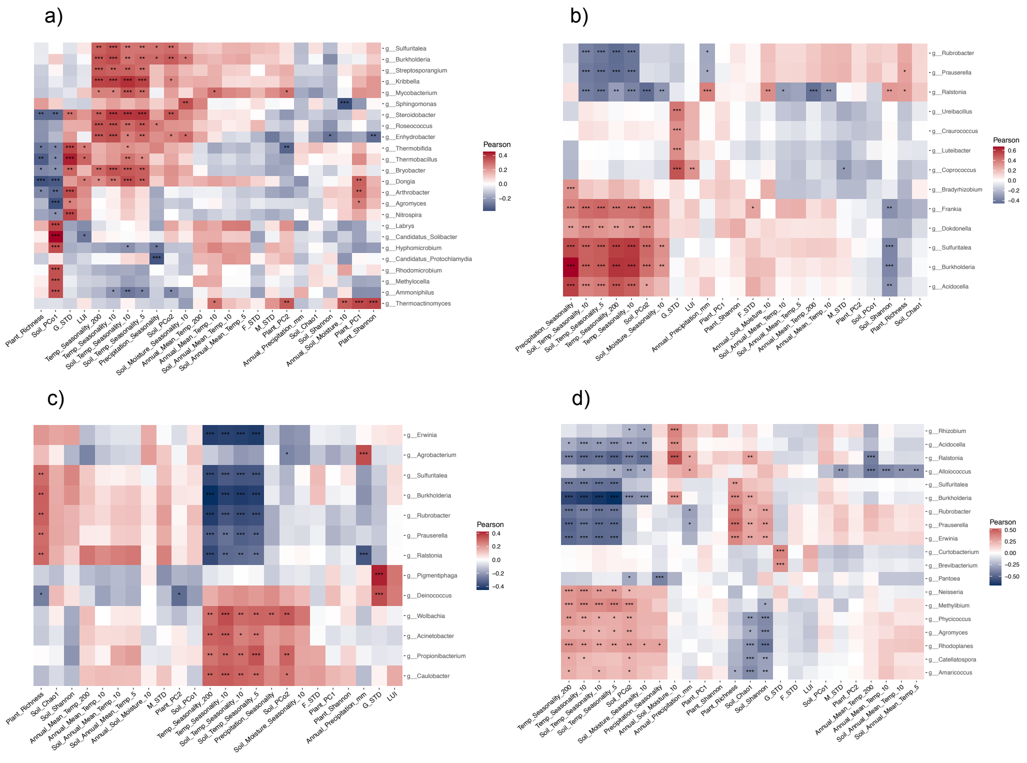

**Supplementary Figure 9.** Heatmap of Pearson correlations between all environmental variables and relative abundances of highly correlated bacterial genera in (a) roots, (b) shoots, (c) flowers, and (d) seeds.

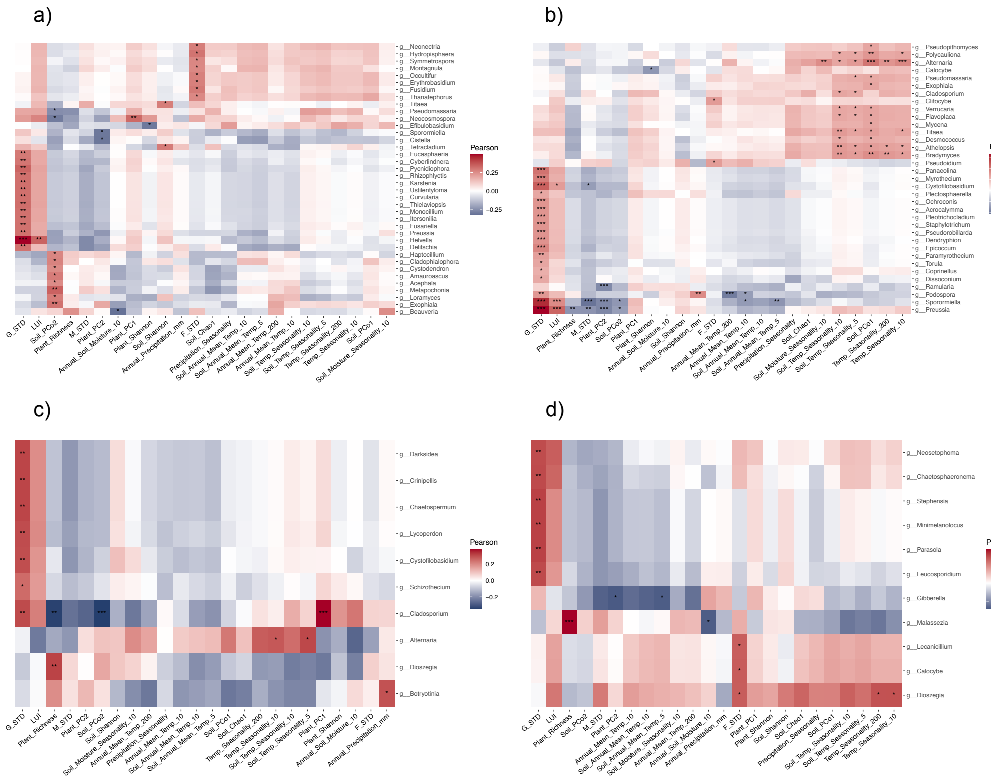


**Supplementary Figure 10.** Heatmap of Pearson correlations between all environmental variables and relative abundances of highly correlated fungal genera in (a) roots, (b) shoots, (c) flowers, and (d) seeds.

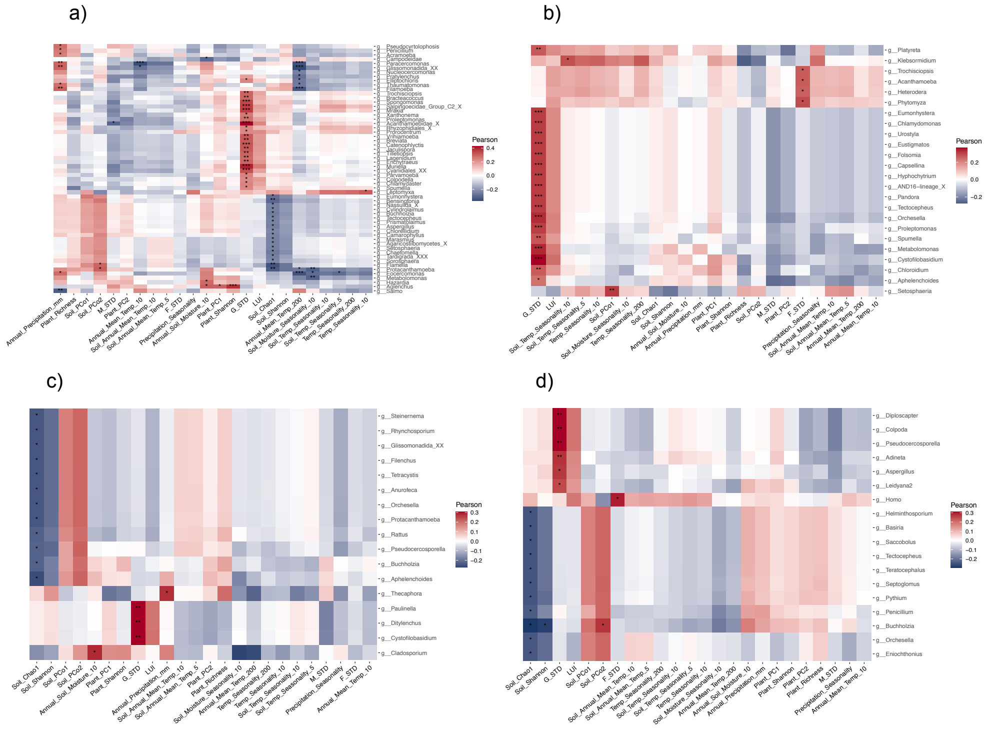


**Supplementary Figure 11.** Heatmap of Pearson correlations between all environmental variables and relative abundances of highly correlated eukaryotic genera in (a) roots, (b) shoots, (c) flowers, and (d) seeds.

**SUPPLEMENTARY TABLES**

**Supplementary Table 1.** Primers and blocking oligos used in this study.

| Primer name | Primer sequence (5'-to-3' orientation) |
|---|---|
| 799F | AACMGGATTAGATACCCKG |
| 1192R | ACGTCATCCCCACCTTCC |
| fITS7 | GTGARTCATCGAATCTTTG |
| ITS4 | TCCTCCGCTTATTGATATGC |
| F1422 | ATAACAGGTCTGTGATGCCC |
| R1797 | TGATCCTTCTGCAGGTTCACCTAC |
| clamp1_BV5_mitoF | GATGAGTGTTCGCCCTTGGTCTACGTGGAT |
| clamp1_BV5_mitoR | CTGCTCAGGGTTCCAAACTCAACGTTGGCA |
| clamp1_ITS2_F | AACCATTAGGTCGAGGGCACGTCTGCCTGG |
| clamp1_ITS2_R | TGAGMGYGGTTACACCACGCATGCGGGTCT |
| clamp9_PV9_F | GATGTATTCAACGAGTCTATAGCCTTGGCC |
| clamp15_PV9_R | TCTCACAACGTCGCAGGCAGCGAACCGCCC |

**Supplementary Table 2.** Contribution of environmental variables to the dbRDA model (vegan:envfit).

a) ROOTS

| | dbRDA1 | dbRDA2 | r2 | Pr(>r) | |
|---|---|---|---|---|---|
| G_STD | 0.21124 | -0.97743 | 0.1169 | 0.001 | *** |
| M_STD | 0.09815 | 0.99517 | 0.0174 | 0.243 | |
| F_STD | 0.99293 | -0.11867 | 0.0094 | 0.451 | |
| LUI | 0.51492 | -0.85724 | 0.1028 | 0.001 | *** |
| Plant_PC1 | 0.47555 | -0.87969 | 0.1156 | 0.001 | *** |
| Plant_PC2 | -0.42413 | 0.9056 | 0.1 | 0.001 | *** |
| Plant_Richness | 0.1337 | 0.99102 | 0.2162 | 0.001 | *** |
| Plant_Shannon | 0.81892 | -0.5739 | 0.0185 | 0.206 | |
| Soil_PCo1 | -0.50667 | 0.86214 | 0.4562 | 0.001 | *** |
| Soil_PCo2 | -0.74359 | -0.66863 | 0.1295 | 0.001 | *** |
| Soil_Chao1 | 0.78725 | 0.61664 | 0.0282 | 0.111 | |
| Soil_Shannon | 0.87638 | 0.48162 | 0.0445 | 0.03 | * |
| Annual_Soil_Moisture_10 | 0.9976 | -0.06925 | 0.1202 | 0.001 | *** |
| Soil_Moisture_Seasonality_10 | -0.78114 | -0.62436 | 0.0627 | 0.009 | ** |
| Soil_Annual_Mean_Temp_10 | -0.91791 | -0.39678 | 0.0146 | 0.279 | |
| Soil_Temp_Seasonality_10 | -0.44233 | -0.89685 | 0.2605 | 0.001 | *** |
| Soil_Annual_Mean_Temp_5 | -0.96703 | -0.25468 | 0.0121 | 0.348 | |
| Soil_Temp_Seasonality_5 | -0.52027 | -0.854 | 0.2504 | 0.001 | *** |
| Annual_Precipitation_mm | 0.86112 | 0.5084 | 0.0677 | 0.002 | ** |
| Precipitation_Seasonality | 0.5805 | -0.81426 | 0.0544 | 0.008 | ** |
| Annual_Mean_Temp_200 | -0.98985 | -0.14212 | 0.1757 | 0.001 | *** |
| Temp_Seasonality_200 | -0.50787 | -0.86143 | 0.1563 | 0.001 | *** |
| Annual_Mean_Temp_10 | -0.98637 | -0.16453 | 0.0753 | 0.002 | ** |
| Temp_Seasonality_10 | -0.45047 | -0.89279 | 0.2141 | 0.001 | *** |

Signif. codes: 0 '***' 0.001 '**' 0.01 '*' 0.05 '.' 0.1 ' ' 1

Permutation: free
Number of permutations: 999

b) SHOOTS

| | dbRDA1 | dbRDA2 | r2 | Pr(>r) | |
|---|---|---|---|---|---|
| G_STD | -0.58521 | -0.81089 | 0.0641 | 0.007 | ** |
| M_STD | 0.36054 | 0.93274 | 0.0681 | 0.006 | ** |
| F_STD | -0.09965 | 0.99502 | 0.0383 | 0.038 | * |
| LUI | -0.9992 | -0.03996 | 0.0041 | 0.747 | |
| Plant_PC1 | -0.715 | 0.69913 | 0.0335 | 0.055 | . |
| Plant_PC2 | 0.76024 | 0.64964 | 0.0132 | 0.329 | |
| Plant_Richness | 0.96952 | -0.24502 | 0.2294 | 0.001 | *** |
| Plant_Shannon | -0.347 | 0.93786 | 0.0724 | 0.002 | ** |
| Soil_PCo1 | 0.97843 | -0.20659 | 0.0093 | 0.472 | |
| Soil_PCo2 | -0.7466 | 0.66527 | 0.3741 | 0.001 | *** |
| Soil_Chao1 | 0.99957 | -0.02929 | 0.0827 | 0.001 | *** |
| Soil_Shannon | 0.74073 | -0.6718 | 0.2178 | 0.001 | *** |
| Annual_Soil_Moisture_10 | 0.74701 | 0.66482 | 0.1556 | 0.001 | *** |
| Soil_Moisture_Seasonality_10 | -0.74603 | 0.66591 | 0.2245 | 0.001 | *** |
| Soil_Annual_Mean_Temp_10 | -0.77495 | 0.63202 | 0.0534 | 0.008 | ** |
| Soil_Temp_Seasonality_10 | -0.80379 | 0.59492 | 0.5583 | 0.001 | *** |
| Soil_Annual_Mean_Temp_5 | -0.68778 | 0.72592 | 0.0553 | 0.009 | ** |
| Soil_Temp_Seasonality_5 | -0.81162 | 0.58418 | 0.5534 | 0.001 | *** |
| Annual_Precipitation_mm | 0.93389 | -0.35755 | 0.094 | 0.001 | *** |
| Precipitation_Seasonality | -0.17916 | 0.98382 | 0.4413 | 0.001 | *** |
| Annual_Mean_Temp_200 | -0.87845 | 0.47784 | 0.1507 | 0.001 | *** |
| Temp_Seasonality_200 | -0.72809 | 0.68548 | 0.503 | 0.001 | *** |
| Annual_Mean_Temp_10 | -0.75412 | 0.65674 | 0.0945 | 0.001 | *** |
| Temp_Seasonality_10 | -0.75467 | 0.65611 | 0.556 | 0.001 | *** |

Signif. codes:  0 ‘***’ 0.001 ‘**’ 0.01 ‘*’ 0.05 ‘.’ 0.1 ‘ ’ 1
Permutation: free
Number of permutations: 999

c) FLOWERS

| | dbRDA1 | dbRDA2 | r2 | Pr(>r) | |
|---|---|---|---|---|---|
| G_STD | 0.69153 | -0.72234 | 0.0869 | 0.001 | *** |
| M_STD | -0.61855 | 0.78575 | 0.1046 | 0.001 | *** |
| F_STD | -0.75279 | -0.65826 | 0.001 | 0.94 | |
| LUI | 0.83544 | -0.54959 | 0.017 | 0.25 | |
| Plant_PC1 | 0.9274 | -0.37406 | 0.0654 | 0.006 | ** |
| Plant_PC2 | -0.50014 | 0.86594 | 0.0772 | 0.001 | *** |
| Plant_Richness | -0.37901 | 0.92539 | 0.0717 | 0.003 | ** |
| Plant_Shannon | 0.99952 | -0.03111 | 0.0116 | 0.38 | |
| Soil_PCo1 | -0.79307 | 0.60912 | 0.1019 | 0.001 | *** |
| Soil_PCo2 | -0.95056 | -0.31053 | 0.2485 | 0.001 | *** |
| Soil_Chao1 | 0.66223 | 0.7493 | 0.0046 | 0.692 | |

| | | | | | |
|---|---|---|---|---|---|
| Soil_Shannon | 0.90306 | 0.42951 | 0.0904 | 0.001 | *** |
| Annual_Soil_Moisture_10 | 0.68503 | -0.72852 | 0.0599 | 0.009 | ** |
| Soil_Moisture_Seasonality_10 | -0.99694 | 0.07823 | 0.1877 | 0.001 | *** |
| Soil_Annual_Mean_Temp_10 | -0.96463 | 0.26362 | 0.0939 | 0.001 | *** |
| Soil_Temp_Seasonality_10 | -0.6913 | -0.72257 | 0.0879 | 0.002 | ** |
| Soil_Annual_Mean_Temp_5 | -0.95162 | 0.30727 | 0.1046 | 0.001 | *** |
| Soil_Temp_Seasonality_5 | -0.80047 | -0.59937 | 0.104 | 0.001 | *** |
| Annual_Precipitation_mm | 0.92017 | -0.39153 | 0.0503 | 0.013 | * |
| Precipitation_Seasonality | -0.85354 | -0.52102 | 0.0853 | 0.001 | *** |
| Annual_Mean_Temp_200 | -0.92559 | 0.37854 | 0.096 | 0.002 | ** |
| Temp_Seasonality_200 | -0.79935 | -0.60087 | 0.1167 | 0.001 | *** |
| Annual_Mean_Temp_10 | -0.8427 | 0.53838 | 0.0607 | 0.004 | ** |
| Temp_Seasonality_10 | -0.82283 | -0.56829 | 0.1463 | 0.001 | *** |

Signif. codes:  0 ‘***’ 0.001 ‘**’ 0.01 ‘*’ 0.05 ‘.’ 0.1 ‘ ’ 1
Permutation: free
Number of permutations: 999

d) SEEDS

| | dbRDA1 | dbRDA2 | r2 | Pr(>r) | |
|---|---|---|---|---|---|
| G_STD | -0.84721 | 0.53125 | 0.0255 | 0.133 | |
| M_STD | 0.9546 | -0.29788 | 0.0143 | 0.32 | |
| F_STD | -0.86137 | 0.50797 | 0.0064 | 0.589 | |
| LUI | -0.45251 | 0.89176 | 0.0098 | 0.458 | |
| Plant_PC1 | -0.19806 | 0.98019 | 0.0603 | 0.005 | ** |
| Plant_PC2 | -0.36462 | 0.93116 | 0.0117 | 0.391 | |
| Plant_Richness | 0.97919 | -0.20294 | 0.1711 | 0.001 | *** |
| Plant_Shannon | 0.11269 | 0.99363 | 0.0624 | 0.004 | ** |
| Soil_PCo1 | 0.82915 | -0.55903 | 0.0204 | 0.166 | |
| Soil_PCo2 | -0.98586 | 0.16756 | 0.3926 | 0.001 | *** |
| Soil_Chao1 | 0.98026 | -0.19773 | 0.1152 | 0.001 | *** |
| Soil_Shannon | 0.91754 | -0.39765 | 0.2109 | 0.001 | *** |
| Annual_Soil_Moisture_10 | 0.67946 | 0.73372 | 0.2564 | 0.001 | *** |
| Soil_Moisture_Seasonality_10 | -0.99465 | -0.10327 | 0.2353 | 0.001 | *** |
| Soil_Annual_Mean_Temp_10 | -0.94176 | 0.33628 | 0.0486 | 0.017 | * |
| Soil_Temp_Seasonality_10 | -0.98997 | 0.14126 | 0.5906 | 0.001 | *** |
| Soil_Annual_Mean_Temp_5 | -0.94977 | 0.31293 | 0.0428 | 0.028 | * |
| Soil_Temp_Seasonality_5 | -0.99579 | 0.09162 | 0.6236 | 0.001 | *** |
| Annual_Precipitation_mm | 0.8461 | 0.53303 | 0.0468 | 0.021 | * |
| Precipitation_Seasonality | -0.61599 | 0.78775 | 0.0972 | 0.001 | *** |
| Annual_Mean_Temp_200 | -0.92024 | -0.39136 | 0.1414 | 0.001 | *** |
| Temp_Seasonality_200 | -0.99023 | 0.13943 | 0.5399 | 0.001 | *** |
| Annual_Mean_Temp_10 | -0.98051 | 0.19647 | 0.0375 | 0.041 | * |
| Temp_Seasonality_10 | -0.99474 | 0.10244 | 0.5907 | 0.001 | *** |

Signif. codes:  0 ‘***’ 0.001 ‘**’ 0.01 ‘*’ 0.05 ‘.’ 0.1 ‘ ’ 1
Permutation: free
Number of permutations: 999

**Supplementary Table 3.** Summary statistics of plant organ microbiome networks generated by network analyzer in Cytoscape 3.10.1.

| Summary Statistics | Roots | Shoots | Flowers | Seeds |
|---|---|---|---|---|
| Number of nodes | 708 | 338 | 119 | 159 |
| Number of edges | 14977 | 4014 | 429 | 1358 |
| Average number of neighbors | 42.308 | 23.751 | 8.286 | 17.082 |
| Network diameter | 6 | 7 | 9 | 7 |
| Network radius | 4 | 4 | 5 | 4 |
| Characteristic path length | 2.522 | 2.794 | 3.718 | 2.924 |
| Clustering coefficient | 0.487 | 0.561 | 0.563 | 0.643 |
| Network density | 0.06 | 0.07 | 0.085 | 0.108 |
| Network heterogeneity | 0.869 | 0.72 | 0.575 | 0.953 |
| Network centralization | 0.191 | 0.162 | 0.102 | 0.23 |
